# Conservation of transcriptional regulatory networks in zebrafish and human periderm facilitates identification of *GRHL1* as an orofacial cleft risk gene

**DOI:** 10.64898/2026.04.27.720742

**Authors:** Sunil K Singh, Annika Helverson, Colin Kenny, Lira Pi, Alexandra Manchel, Sarah W. Curtis, Kelsey Robinson, Kaylia M Duncan-Field, Edward B Li, Eric C Liao, Justin Cotney, Elizabeth J Leslie-Clarkson, Patrick Breheny, Robert A Cornell

## Abstract

Most heritable risk for orofacial clefts (OFC) remains unassigned to specific genes or loci. *IRF6* and *GRHL3*, two established OFC risk genes, encode transcription factors (TFs) essential for the differentiation of the periderm, a transient embryonic tissue required for secondary palate fusion. To identify novel risk candidates, we modeled the zebrafish periderm transcriptional regulatory network (TRN). Using single-cell multiome sequencing (RNA-seq and ATAC-seq) from shield-stage embryos, we inferred TF-to-target gene connections by integrating correlated gene expression with TF binding site predictions within chromatin elements open in periderm cells. We generated sets of gold-standard edges by conducting RNA-seq on TF-depleted embryos and ChIP-seq/CUT&RUN on wild-type embryos and used them to benchmark model performance. Within the top-performing model, zebrafish periderm modules are strongly preserved in human embryonic periderm and orthologs of human OFC-associated genes have higher centrality and edge-sum scores than non-associated genes. Functional validation confirmed the network’s predictive power: depleting high-centrality TFs, including *grhl1*, *klf6a*, *tead3b*, and *klf17*, disrupted periderm differentiation in sensitized embryos. Moreover, analysis of whole-genome sequencing data from 2,415 OFC trios identified 15 individuals with rare or *de novo GRHL1* variants, four of which introduced premature stop codons. This study establishes *GRHL1* as a novel OFC risk gene and highlights the power of cross-species gene regulatory network analysis to prioritize candidates for rare variants in complex structural birth defects.

## Introduction

The role of rare genetic variants in pathogenesis of orofacial cleft (OFC), a congenital birth defect which comprises cleft lip, cleft palate, or a combination of the two, is incompletely understood. In most of the cases, OFC occurs as the only anomaly and is termed non-syndromic (nsOFC)^1^. Evidence for a genetic contribution to the etiology of nsOFC is that the concordance of nsOFC in monozygotic twins (about 50%) is higher than that in dizygotic twins (about 5%) ^2,3^. Multiple genome wide association studies (GWAS) and meta-analyses of them have identified more than 40 loci where common variants are associated with risk for OFC ^4–10^. There is debate regarding how to accurately infer the fraction of overall heritability of complex diseases ^11^. However, most GWAS have limited power to evaluate polymorphisms with minor allele frequencies under 5% ^12^ and it is well accepted that common variants contribute only a fraction of the heritable risk for nsOFC ^13,14^. Rarer variants can be identified through exome or whole genome sequencing of individuals with OFCs, but such efforts identify thousands of rare variants in every individual, not just those with a birth anomaly ^15–17^. Moreover, all individuals harbor many variants that computer algorithms predict alter protein function and many more that have unknown functional significance ^18^. It is thus an outstanding challenge in the field to identify the rare variants that contribute risk for OFC.

To prioritize genes for their likelihood of harboring variants that will cause OFC pathogenesis it is helpful to know the genes that govern differentiation of craniofacial tissues. Among these tissues is periderm, a simple squamous epithelium present in embryos prior to the maturation of epidermis that serves as barrier to dehydration ^19^. In murine embryos periderm is generated after stratification of early embryonic epidermis and oral epithelium ^20,21^. The periderm prevents interepithelial adhesions among limbs, oral structures and digits ^20,22^; paradoxically, it facilitates the transient fusion of digits, eyes, and pinnae in early post-natal mice ^23,24^. Interestingly, multiple genes that are necessary differentiation of periderm, including *IRF6*, *GRHL3*, and *TP63,* when mutated cause syndromes with OFC-risk genes ^25–27^. In addition, atypical protein kinase C (aPKC) activity is essential for periderm differentiation in frogs and fish ^28,29^, and *PRKCI*, encoding an aPKC, is also an OFC risk gene ^29^. Therefore, other genes that participate in periderm differentiation are candidates to harbor mutations that contribute risk for OFC.

In zebrafish embryos shortly before gastrulation superficial blastomeres differentiate into enveloping layer (EVL), a simple squamous epithelial monolayer ^30,31^. The EVL forms a separate lineage and becomes the periderm that surrounds the embryos after gastrulation ^31,32^. Differentiation of zebrafish EVL depends on many of the same transcription factors (TFs) as that of mammalian periderm. For instance, in *irf6* maternal zebrafish mutants, RNA-seq of reveals strongly reduced expression of virtually all genes whose expression normally characterizes the EVL ^33^; such mutants, or embryos expressing a dominant negative form of Irf6, never initiate epiboly and die at about 6 hpf ^34,35^. Zygotic *grhl3* loss-of-function mutants complete epiboly but rupture at about 11 hours post fertilization ^36^; housing conditions can prevent this rupture and mutants that survive to larval stages have constricted oral cavities and intestines, similar to mouse *irf6* mutants ^37^. Maternal *ikk1* mutants undergo delayed epiboly and frequently rupture during epiboly and have reduced expression of periderm markers ^38^. And, as mentioned above EVL differentiation also fails in embryos treated with an inhibitor of atypical PKCs, which include the OFC-associated *PRKCI* ^29^. In summary, many of the same regulatory molecules regulated differentiation of zebrafish EVL and of mammalian oral periderm. Because zebrafish embryos develop externally, EVL is an easily accessible model to investigate potential drivers of oral periderm differentiation.

To identify novel regulators of EVL differentiation we sought to construct models of the EVL transcriptional regulatory network (TRN). TRNs are comprised of the set of transcription factors (TFs) expressed in a given tissue and their regulatory connections (edges) to target genes; they are a reductionist description of the genetic circuitry that governs cellular differentiation ^39–41^. TRNs are cumbersome to deduce experimentally but can be inferred computationally ^42^. TRN-predicting algorithms infer edges between genes if their expression is correlated in multiple gene expression profiles (gene expression co-variation). Single cell RNA sequencing experiments, which can yield hundreds of profiles for a given cell type, are a useful source for these datasets, with some caveats, including dropouts ^43^. Some algorithms additionally infer edges between TFs and target genes if the canonical TF binding site (TFBS) is present in open chromatin near the transcriptional start site of the target gene; we will refer to this type of evidence as “prior” following the nomenclature of the study that inspired this one ^44^. Single cell ATAC seq data can be useful in this context as they can reveal cell-type specific open chromatin elements; however, they require large numbers of cells per cluster to overcome the sparseness inherent in there being just two copies of each genomic locus in every cell ^45^.

We predicted that OFC-associated TFs are relatively centrally located in the EVL TRN, and reasoned that if this prediction is correct, then other centrally located TFs will be candidates to harbor variants that cause OFC. We use single cell multiomic data from zebrafish embryos at the beginning of epiboly and four TRN-inferring algorithms to generate multiple models of the EVL TRN. We generate sets of gold-standard edges and identify the TRN model that best predicts these edges and use the best performing model to rank the centrality of TFs. Finally, we knock down expression of TFs predicted to be highly central in the EVL TRN and in such embryos assess the integrity of periderm differentiation.

## Material and methods

### Zebrafish

All the procedures involving zebrafish were approved by the Institutional Animal Care and Use Committee (IACUC) at the University of Iowa and University of Washington, Seattle and follow NIH guidelines. Zebrafish wild-type AB strain embryos were used for all the experiments unless specified otherwise. Zebrafish embryos were grown in embryo media (5.03 mM NaCl, 0.17 mM KCl, 0.33 mM CaCl_2_, 0.33 mM MgSO_4_, 0.1% Methylene blue) at 28.5°C and used for procedures at different stages according to the standard staging protocol ^31^.

### Single cell multiome (ATAC + RNA-seq)

So that we could use the resulting data in a gene-knockdown study that is in preparation, we first injected wild-type zebrafish embryos at the one-to-four cell stage with ∼5 ng of the standard negative control morpholino from GeneTools (Philomath, Oregon). At 6 hpf (shield-stage) we manually dechorionated two groups of 100 embryos and deyolked them using ice-cold phosphate buffer saline solution (1x PBS). We dissociated the embryos with 0.25% Trypsin for 5 minutes at 37 °C and then collected cells by spinning in a refrigerated centrifuge for 2 minutes at 400 rcf. Cell suspensions were processed with 10X Genomics Nuclei isolation kit to get single nuclei suspension. Nuclei were subjected to transposition and adapter ligation in transposition mix. Finally, approximately 12,000 cells per replicate were loaded on 10x Genomics Chromium using Next-GEM standard protocol. ATAC and gene expression libraries were prepared and sequenced on Illumina HiSeq 2500 sequencer. Raw sequence data was mapped on zebrafish danRer11 genome using 10X Genomics Cell Ranger ARC software (https://www.10xgenomics.com/support/software/cell-ranger/latest). Mapped fragment files were analyzed using Seurat package (Version 4.0; ^46^). Cell clusters were identified using the top 5 most specifically expressing genes in each cluster. Open chromatin regions were identified using MACS2 ^47^.

### Prior information matrix

EVL cluster was identified based on expression of EVL-specific marker genes (*eppk1, krt4*, *krt8*, *spaca4l*, and *krt18a*). The PWMs were downloaded from catalog of inferred sequence binding preferences (cis-bp; http://cisbp.ccbr.utoronto.ca/) and swissRegulon https://swissregulon.unibas.ch/pages/) databases (**Table S1**). For prior information matrix, chromatin regions accessible in EVL cells were searched for occurrences of TF PWM using the FIMO tool ^48^. The regions with occurrences of a particular TF’s PWM were annotated to all the genes (TSS) within 100 kb of that region. A binary prior information matrix was generated based on presence (denoted as “1”) or absence (denoted as “0”) of a TF PWM in the open chromatin region nearby a gene.

### Network inference

A list target genes TRN inference was prepared comprising of 3,787 genes that were expressed in 10 percent or more of the EVL cells regardless of their relative expression in EVL cells versus in other cell types. Gene expression matrix was prepared by taking the FPKM values from gene expression (in rows) data of 394 EVL cells (in columns). Very low expressing genes (expressed in less than 10 percent cells) were excluded from the matrix.

Gene expression and prior information matrices were used to infer TRN models using four different algorithms: WGCNA ^49^, GENIE3 ^50^, gLASSO (graphical lasso) ^51^, and Inferelator 3.0 ^52^. All four algorithms produce a score per edge (correlation, regression coefficient, etc.) that reflects the strength of evidence for that edge. For WGCNA and GENIE3.0, these scores were upweighted by a factor that depends on the prior strength (0 meaning no change, 1 meaning that the score is doubled, and so on). For gLASSO and INFERELATOR, the strength was directly represented by a penalty or prior, respectively.

### Zebrafish embryo micro-injections

Zebrafish embryos were injected at 1-2 cell stage with approximately 3nL of 1 mg/mL (1.5 mg/mL for *grhl3* MO) morpholino **(Table S2**; GeneTools Inc.) solutions individually. Injected embryos were kept in 100 mm tissue culture dishes with embryo medium (5.03 mM NaCl, 0.17 mM KCl, 0.33 mM CaCl_2_, 0.33 mM MgSO_4_, 0.1% Methylene blue) at 28.5°C until harvested for different assays. Approximately 120 embryos were injected for each group. Injection experiments were independently replicated on at least three days. Unfertilized and dead embryos were removed from all the dishes at 4 hpf. Standard control MO (GeneTools Inc.) was used as negative control for the injections.

### CRISPR-Cas9

Custom Alt-R^TM^ CRISPR-Cas9 guide RNAs (**Table S3**) were designed using Integrated DNA Technologies (IDT, Inc.) design tool (https://www.idtdna.com/site/order/designtool/index/CRISPR_CUSTOM). The CRISPR guide RNAs (crRNAs), tracrRNA, and Cas9 protein were ordered from IDT, Inc. and RNPs were generated using previously reported protocol ^53^. Briefly, crRNA:tracrRNA duplex stock solutions (50μM) were made by mixing 100 uL each of individual crRNAs with tracerRNA. Finally, we mixed 1ul of 25μM crRNA:tracrRNA duplex with 1 ul of 25μM Cas9 stock, 2 ul H_2_O, and 1ul 0.25% phenol red solution to prepare 5μM crRNA:tracrRNA:Cas9 RNP solutions for micro-injections. This solution was incubated for 5 minutes at 37°C and then placed at room temperature until injected into the zebrafish embryos at 1-2 cell stage (∼3 nL per embryo). Approximately 120 embryos were injected for each group. Injection experiments were independently replicated on at least three days. Unfertilized and dead embryos were removed from all the dishes at 4 hpf. The crispr guide against zebrafish Tyrosine (*tyr*) gene was used as negative control.

### Gold standard data sets

Total RNA samples were extracted from MO injected zebrafish embryos (40 embryos per replicate) using RNAqueous™ Total RNA Isolation Kit (Cat# AM1912; Thermo Fisher Scientific) at approximately 5.5hpf following the standard protocol. The RNA samples were checked for quality and quantity using Qubit 3.0 fluorometer and NanoDrop 2000 spectrophotometer (Thermo Fisher Scientific). Approximately 100 ng of total RNA samples were sent to Novogene, Inc. for mRNA sequencing. RNA-seq library preparation was performed using standard protocol and libraries were sequenced on Illumina HiSeq 2500 sequencing system. The resulting raw reads were quality filtered using trimGalore (https://github.com/FelixKrueger/TrimGalore) and aligned to the zebrafish (danRer11) reference genome by employing STAR RNA-seq aligner ^54^. Reads mapped to specific genes were quantified using featureCounts tool from Subread package ^55^. DESeq2 ^56^ and ashr ^57^ packages were used to identify the differentially expressed genes (DEGs). Gold standard true positives were restricted to target genes with a local false sign rate (lfsr) of 0.01 or lower.

For Irf6 ChIP-sequencing, maternal-null *irf6*^-/-^ embryos ^34^ were injected with zebrafish *irf6* 3xFLAG-tagged mRNA at the 1-cell stage, raised to 4-5 hpf and used for ChIP-seq with an anti-FLAG M2 antibody as described earlier ^58^. CUT&RUN assays for Grhl3 and Tfap2a (2 replicates each) were performed on lysates of 6 hpf zebrafish embryos using custom made anti-Grhl3 antibody and anti-Tfap2a antibody (Cat# PA5-17359; Thermo Fisher Scientific). ChIP and CUT&RUN libraries were prepared, sequenced, and quality filtered as described above. ChIP-seq and CUT&RUN sequencing reads were aligned to zebrafish danRer11 reference genome using Bowtie 2 aligner ^59^ with standard parameters. TF binding peaks were called using MACS2 tool ^47^. For gold standard chromatin datasets, we annotated called peaks to the nearest genes.

### Validation of network predictions

Applying a threshold to each algorithm’s scores, the edge predictions can be compared to the ground truth to obtain precision (percent of edges above threshold that are true positives) and recall (percent of true positives above threshold). Varying the threshold produces a precision-recall curve. We used eigenvector centrality, which accommodates continuous edge scores, to quantify the importance of TFs as central players in the network. For non-TF genes, which always have low centrality, we quantified their importance by the weighted sum of edges pointing to it across all TFs (“EdgeSum”). The Walktrap community detection algorithm ^60^ was used to cluster TFs in the network into modules.

### Enrichment of OFC-associated genes in predicted network

To identify the OFC-associated TFs and non-TF (effector) genes in our network model, we curated a list of genes by using OFC gene panels from NIH (USA), NHS (UK), and previously published reports (**Table S13**). Zebrafish orthologs of these OFC associated genes were identified using ZFIN database (zfin.org/). Wilcoxon rank-sum tests were used to compare the centrality and EdgeSum scores of TFs and effector genes, classified by whether or not they appear in the gene panel.

### Network preservation analysis

For cross-specifies comparison between zebrafish and human, we utilized single nucleus RNA-seq data and cell subtype annotations from human craniofacial tissue during embryogenesis. Genes from the zebrafish WGCNA EVL Module 1 were subset to include only those with one-to-one orthology between zebrafish and human. These genes were then used to calculate a module score for EVL Module 1 using the “AddModuleScore” function in Seurat. The median module scores were then ranked across cell subtypes and plotted according to their rank.

We used the R package DenovolyzeR v. 0.2.0 ^61^ to test enrichment of de novo variants (DNs) in a dataset of OFC case-parent trios (Gabriella Miller Kids First Pediatric Research Consortium). Enrichment is calculated by comparing the expected number of variants, as determined by mutation models described by Samocha ^62^, to the observed number of variants in a given gene or group of genes using the ‘DenovolyzeByClass’ and ‘includeGenes’ functions. We first compared the genes with DNs in trios with OFCs to those with calculated mutational rates in the R package ‘DenovolyzeR’ using the ‘viewProbabilityTable()’ function. We then tested enrichment of the three WGCNA module gene lists, subset to those with one-to-one orthology between zebrafish and human, across all OFC trios.

### Whole genome sequencing (WGS) and quality control (QC)

The cohort for this study came from three different sequencing initiatives: a study by the Gabriella Miller Kids First Pediatric Research Program (GMKF) with participants with any orofacial cleft (OFC)(N= 3,570), a study by GMKF focusing on participants with cleft lip only (CL)(N=1,892), and another study focusing on participants with cleft palate only (CP)(N=1,625). The combined cohort consisted of 2,415 individuals with any type of orofacial cleft (OFC), of which 2,112 belong to complete trios. There were 798 cleft lip (CL), 1,062 cleft lip and palate (CLP), and 554 cleft palate (CP) probands. This is a multi-ancestry cohort with participants recruited from the United States, Argentina, Turkey, Hungary, Spain, Colombia, the Philippines, and Taiwan. Participant recruitment was done at regional treatment or research centers after review and approval by each site’s institutional review board (IRB) and the IRB of the affiliated US institutions (e.g., the University of Iowa, the University of Pittsburgh, and Johns Hopkins University). Participants were assessed at the time of recruitment for additional clinical features and are largely classified as having an isolated cleft, but participants with additional clinical features consistent with a syndromic diagnosis were not excluded.

Samples from both GMKF Program samples were sequenced at the McDonnell Genome Institute (MGI), Washington University School of Medicine in St. Louis, MO, or the Broad Institute in Boston, MA. The variant calling and quality control methodology for the 1,190 GMKF trios has been described in detail previously ^10,63,64^, and the quality control for the 1,892 participants from the CL study was done similarly. For the 1,625 participants that were sequenced as part of the CP study, whole genome sequencing was performed at the Center for Inherited Disease Research (CIDR) at Johns Hopkins University. The variant calling and quality control methodology has been described in detail previously ^65^.

### Variant annotation and filtering

From the WGS data, we extracted all variants in *GRHL1*, *KLF6*, *KLF17*, and *TEAD3* using MANE select transcripts for GRHL1 (NM_198182.3 and ENST00000324907.14), KLF6 (NM_001300.6 and ENST00000497571.6), KLF17 (NM_173484.4 and ENST00000372299.4), and TEAD3 (NM_001256037.3 and ENST00000338863.13). Variants were annotated using Variant Effect Predictor (VEP) ^66^ with Ensembl release 115. We then filtered variants based on consequence and minor allele frequency (MAF) in gnomAD v.4.1 ^67^. Variants were removed if they were observed with MAF >0.1% in any population in either exomes or genomes, and if they had non-coding or non-protein-altering consequences (i.e., synonymous variants without predicted splicing effects were removed). Remaining variants were then evaluated for potential pathogenicity based on predicted consequences, CADD score (≥25 considered predicted damaging), and inheritance pattern. Details of variants detected are in Table 26.

### Statistical testing

Using the set of complete OFC trios (n=688 CL, n=950 CLP, n=474 CP), we generated transmission counts for each qualifying variants (MAF<0.1%, predicted protein-altering consequence, CADD >25) using the transmission disequilibrium test (TDT) function in PLINK 1.9 ^68^. We then performed an exact binomial test to evaluate *GRHL1* for a burden of rare variant transmission. Results were considered significant if p<0.05.

## Results

### Extracting sets of genes expressed and chromatin regions open in EVL cells from single cell multiome (RNA and ATAC-seq) data

The design of this project is summarized in **Figure 1**. We conducted 10X genomics single-cell (sc) RNA-seq and assay-for-transposon-accessible-chromatin (ATAC)-seq (sc-Multiome) assay on wild-type zebrafish embryos at 6 hours post fertilization (hpf, shield-stage). We aggregated sequencing data from two replicates. We generated uniform-manifold-approximation-and-projection (UMAP) graphs based on a combination of sc-RNA-seq data and sc-ATAC-seq data datasets (weighted-nearest neighbor or WNN analysis) (**Figure 2A**). Using the top five most-highly expressed genes in each cluster we predicted the cell type corresponding to many of the clusters (**Figure S1**; **Table S4**). The EVL cluster, which contained 394 cells, was readily identified as it contained cells expressing levels of *eppk1, krt4* (**Figure 2B**), *krt8*, *spaca4l*, and *krt18a*.

**Figure 1:**
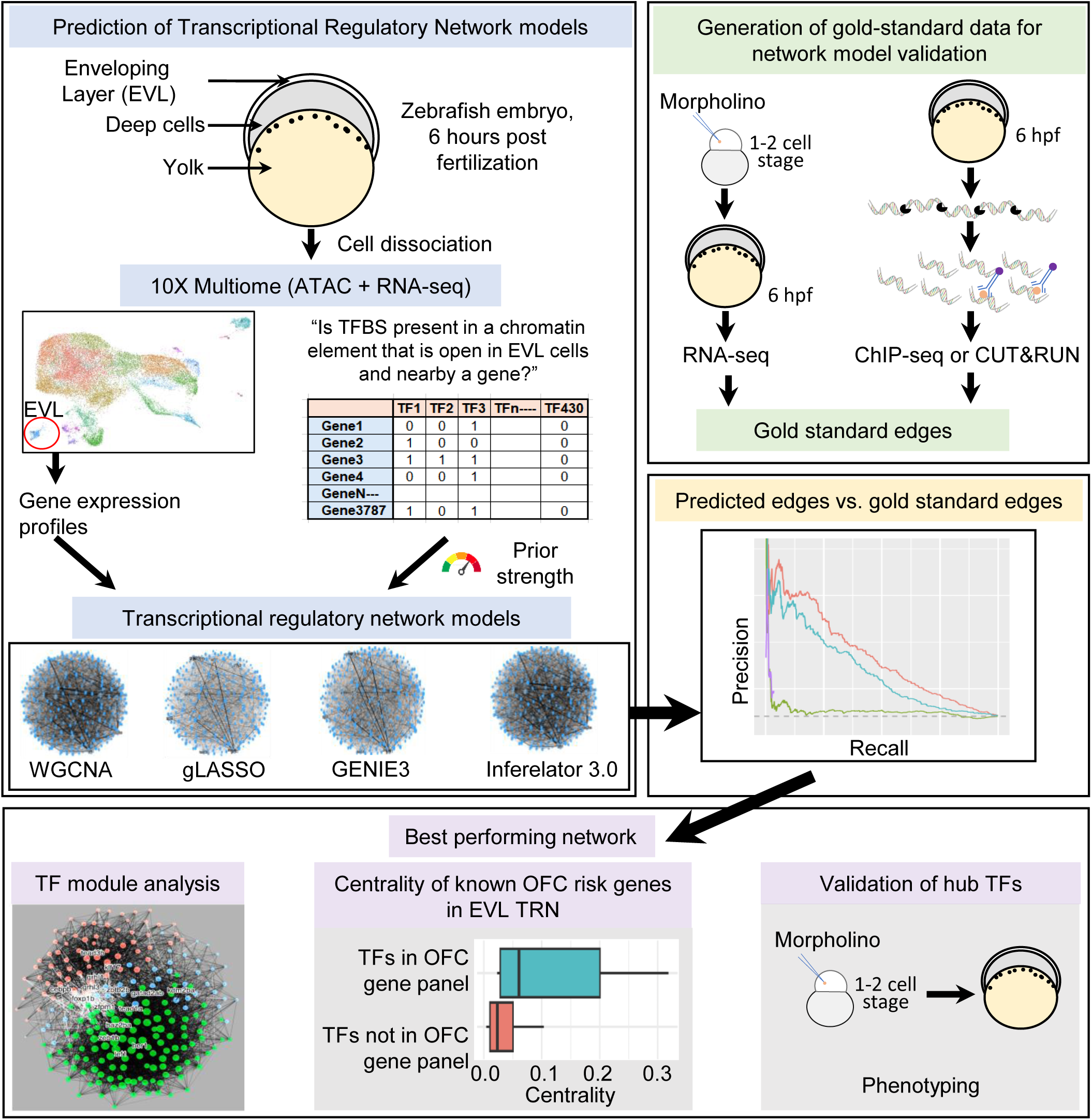
Schematic summary of workflow in this project.

**Figure 2:**
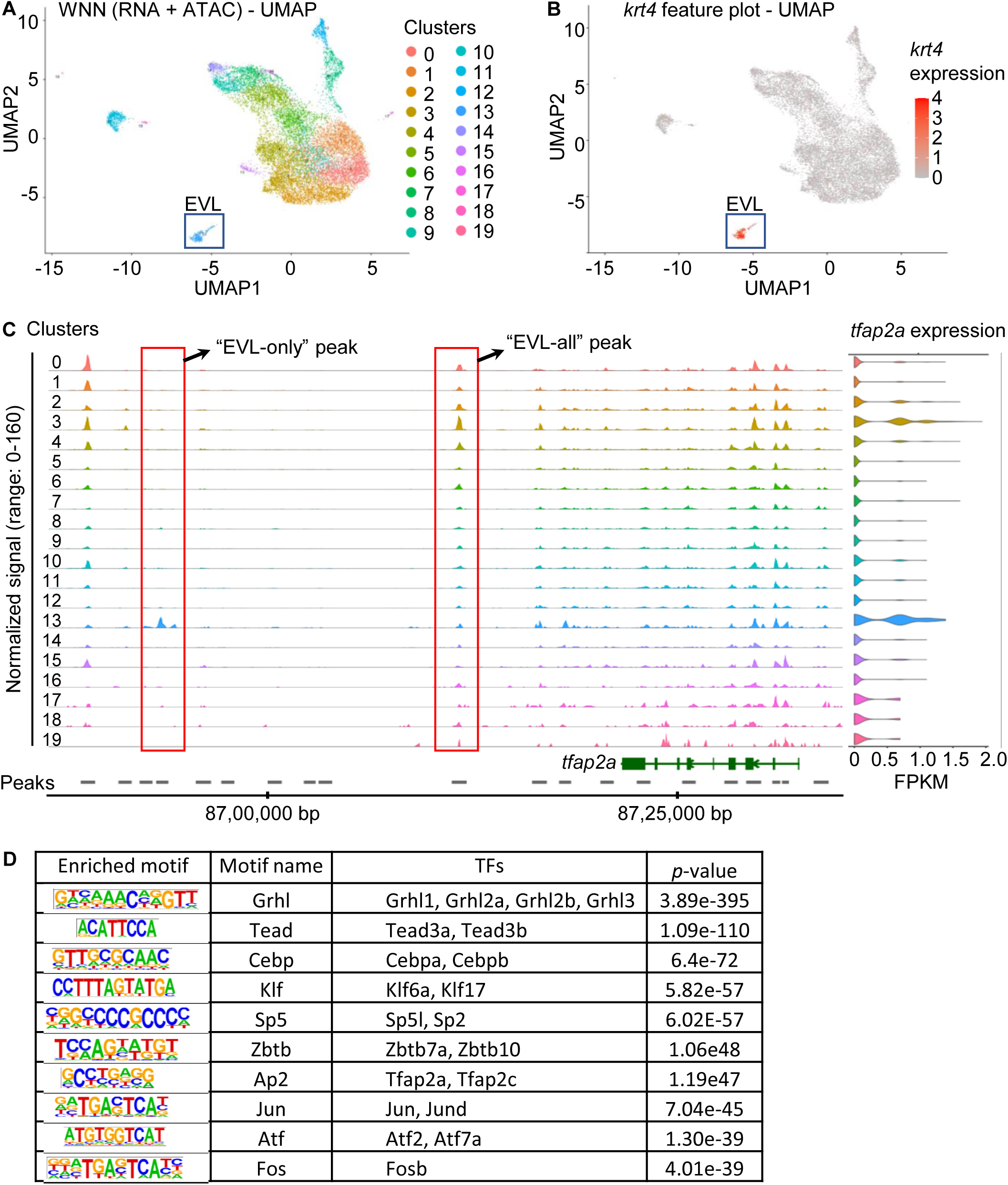
Single cell multiome (ATAC + RNA) sequencing on shield stage zebrafish embryos. A) Single-cell multi-omics data were collected from zebrafish embryos at 6 hours post fertilization using 10x Genomics technology. Uniform Manifold Approximation and Projection (UMAP) visualization of single cells, with each cell colored according to its cluster; the UMAP coordinates are based on combined RNA and ATAC (WNN) data. B) UMAP visualization of single cells at 6-somite stage cell clusters colored by expression level of *krt4* (EVL cell marker) gene. C) Genome browser track showing chromatin accessible peaks for the indicated clusters in genomic region around *tfap2a* gene. D) List of top10 enriched TF-binding motifs in EVL-all peaks with their corresponding p-values and annotations.

We first identified the set of genes to include in the EVL TRN models (i.e., the *target gene list*). We included the 3,787 genes that were expressed in 10 percent or more of the EVL cells regardless of their relative expression in EVL cells versus in other cell types (**Table S5**). This list included 226 genes encoding transcription factors (TFs). In the TRN models, all edges are directed and emanate from TFs to other genes in the list, including those encoding TFs.

We next created a *prior matrix* of transcription factors from the target gene list versus all genes in the list, reflecting whether for each TF a sequence motif corresponding to its predicted binding site (i.e., TFBS) is present in one or more EVL regulatory element near the transcription start site of each target gene. We queried the sc-ATAC-seq data and identified chromatin elements that were accessible solely in EVL cells (*EVL-only* ATAC-seq peaks) or in many or all cell types (*EVL-all* ATAC-seq peaks) ^69^ (**Figure 2C**; **Table S6**). The TFBS for about two thirds of the TFs expressed in EVL (i.e., 151 TFs) were present in catalogs of inferred sequence binding preferences (cis-bp, http://cisbp.ccbr.utoronto.ca/; or SwissRegulon, https://swissregulon.unibas.ch/pages/) (**Table S1**). We used FIMO ^48^ to find occurrences of these TFBS within the EVL regulatory elements (i.e., EVL-all ATAC-seq peaks). Such elements were enriched for the TFBS of many known or proposed regulators of periderm differentiation, including the Grhl, Klf, Tead, and Tfap2 families (**Figure 2D**). We assigned open chromatin elements to genes whose transcription start sites were within 100 kb (**Table S7**).

We used the expression profiles from 394 EVL cells (**Table S8**) and the prior matrix as input to four algorithms that generate TRN models: WGCNA ^49^, GENIE3 ^50^, gLASSO (graphical lasso) ^51^, and Inferelator 3.0 ^52^. All these algorithms utilize co-variance of gene expression in multiple expression profiles as the basis of network edges; gLASSO and Inferelator 3.0 adjust the results according to the prior matrix. WGCNA and GENIE3 do not inherently factor in prior evidence but we applied a post-analysis weighting step to their results so that TF-to-target gene edges lacking support in the prior matrix were penalized (Methods). In each algorithm we varied the weight of the prior-based penalty from zero to four towards yielding network models of optimal performance (assessed below). The gLASSO method has an additional tuning parameter called *alpha* that controls the relative contributions of lasso and ridge penalties in the elastic net model ^70^. With alpha set to zero, gLASSO is effectively identical to WGCNA, whereas with alpha set to one, there is no ridge penalty, and it functions similarly to Inferelator 3.0. Applying each of the four algorithms over a range of these tuning parameters yielded more than 50 models of the EVL TRN (**Tables S9 – S12**).

### Generation of gold standard datasets of TF-to-target-gene edges

To determine which model of the EVL TRN most accurately predicted true TF-to-target-gene edges, we generated gold-standard datasets of such edges. We carried out bulk RNA-seq on embryos depleted of expression of *irf6* **(Figure 3A)**, *grhl3* **(Figure 3B)**, *tfap2a* **(Figure 3C),** *grhl1* **(Figure 3D)**, *klf17* **(Figure 3E)**, *cebpb* **(Figure 3F)**, *tead3b* **(Figure 3G)**, *klf6a* **(Figure 3H)**, *gata3* **(Figure 3I)**, *zbtb10* **(Figure 3J)**, and *znf750* ^71^, either by mutation (*irf6*) or by injecting anti-sense morpholino oligonucleotide (MO) (all the others). We chose these TFs because a) they are all expressed in the EVL at 5.5 hpf ^72^, b) in many cases their binding sites are enriched in chromatin regions open in zebrafish periderm cells at 11 hpf, inferred from bulk ATAC-seq ^73^, and c) because they emerged as key TFs in the initial phase of TRN analysis. To evaluate the efficacy of the splice blocking MOs we examined the RNA seq profiles at the intended splice site target - in each case the MO affected splicing of the intended transcript (**Figure S2**). We compared expression profiles from TF-depleted embryos versus control embryos and identified differentially expressed genes (DEGs) for each TF using the DESeq2 ^56^ and ashr ^57^ packages (**Tables S13 - S22**). We used the sets of DEGs as *Knock-down (KD) RNA-seq* gold standard datasets.

**Figure 3:**
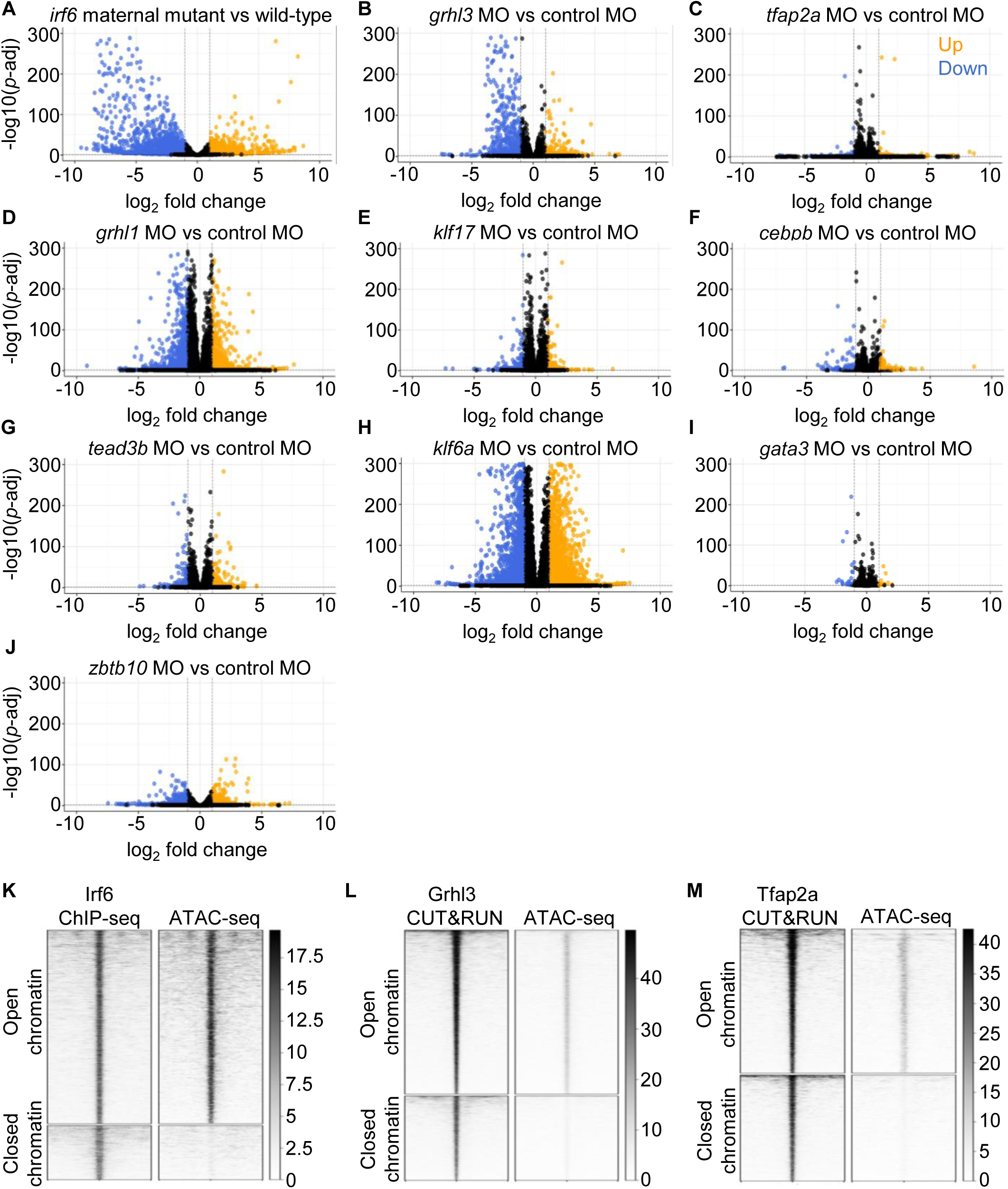
Gold-standard data for comparing the performance of TRN models. A) Volcano plot showing DEGs identified in RNA-seq on *irf6* maternal mutant embryos vs wild-type embryos. Volcano plots showing DEGs identified in RNA-seq on B) *grhl3* MO, C) *tfap2a* MO, D) *grhl1* MO, E) *klf17* MO, F) *cebpb* MO, G) *tead3b* MO, H) *klf6a* MO, I) *gata3* MO, and J) *zbtb10* MO vs standard control MO injected embryos at 5.5 hpf. Heatmaps showing K) Irf6, L) Grhl3, M) Tfap2a binding peaks in open and closed chromatin regions (Shield-stage ATAC-seq data: NCBI GEO - GSM2837497 and GSM2837498).

Because differential expression alone cannot distinguish between direct and indirect regulatory interactions, we also performed ChIP-seq or CUT&RUN assays to identify targets directly regulated by Irf6, Tfap2a or Grhl3. We conducted anti-Myc ChIP-seq on lysates of 6 hpf embryos injected with mRNA encoding Myc-epitope-tagged Irf6 (one replicate; **Table S23**; **Figure 3K)**; we also conducted anti-Grhl3 CUT&RUN (**Table S24; Figure 3L**) and anti-Tfap2a CUT&RUN (**Table S25; Figure 3M**) on lysates of uninjected 6 hpf embryos (two replicates each). The majority of Grhl3 and Tfap2a peaks were in intergenic regions while Irf6 peaks were predominantly in promoter regions (**Figure S3**); anti-IRF6 ChIP-seq peaks detected in human keratinocytes also had this quality ^74,75^. If a ChIP-seq or CUT&RUN peak of a given TF was a) in open chromatin (from an ATAC-seq analysis of zebrafish embryo lysates ^76^) and b) within 100 kb of the transcription start site of a DEG for that TF we considered the gene to be directly regulated by the TF. We referred to these three sets of genes as *KD RNA-seq + TF peaks* gold-standard datasets.

To determine the algorithm and set of parameters that yielded the most accurate model of the EVL TRN across TFs we plotted comparisons of TF-to-target-gene edges in the models to those in the gold-standard datasets. The partial area under the precision-recall curve (pAUPRC) refers to the area under the curve above a line representing random guesses (dotted lines in **Figure 4A-D**) and thus conveys the ability of a model to predict gold-standard edges, which is the model’s performance. The WGCNA algorithm yielded either the best pAUPRC, or one tied for best, for the three *KD RNA-seq + TF* peak datasets (**Figure 4E**). For the Grhl3 and Irf6 edges, a minimal value of prior gave the best pAUPRC, but for Tfap2a edges, a maximal one did. In hopes of finding a stronger consensus for the optimal value of prior we next evaluated the ability of the models to predict edges in the *KD RNA-seq* datasets. For six of the eight *KD RNA-seq* datasets, WGCNA again yielded the best results (**Figure 4D**). Using WGCNA, for one of these datasets, Znf750, a maximum value for the prior weight yielded a small benefit, but for the rest a prior weight of value of zero or one half performed optimally (**Figure 4E**). Merging the results, the EVL TRN model generated by the WGCNA algorithm and a prior weight of 0.5 yielded the best pAUPRCs across gold standard edges, and therefore we picked this model for upcoming analyses. We note that in a previous study, there was similarly no consensus across transcription factors for an optimal value of prior weight ^44^.

**Figure 4:**
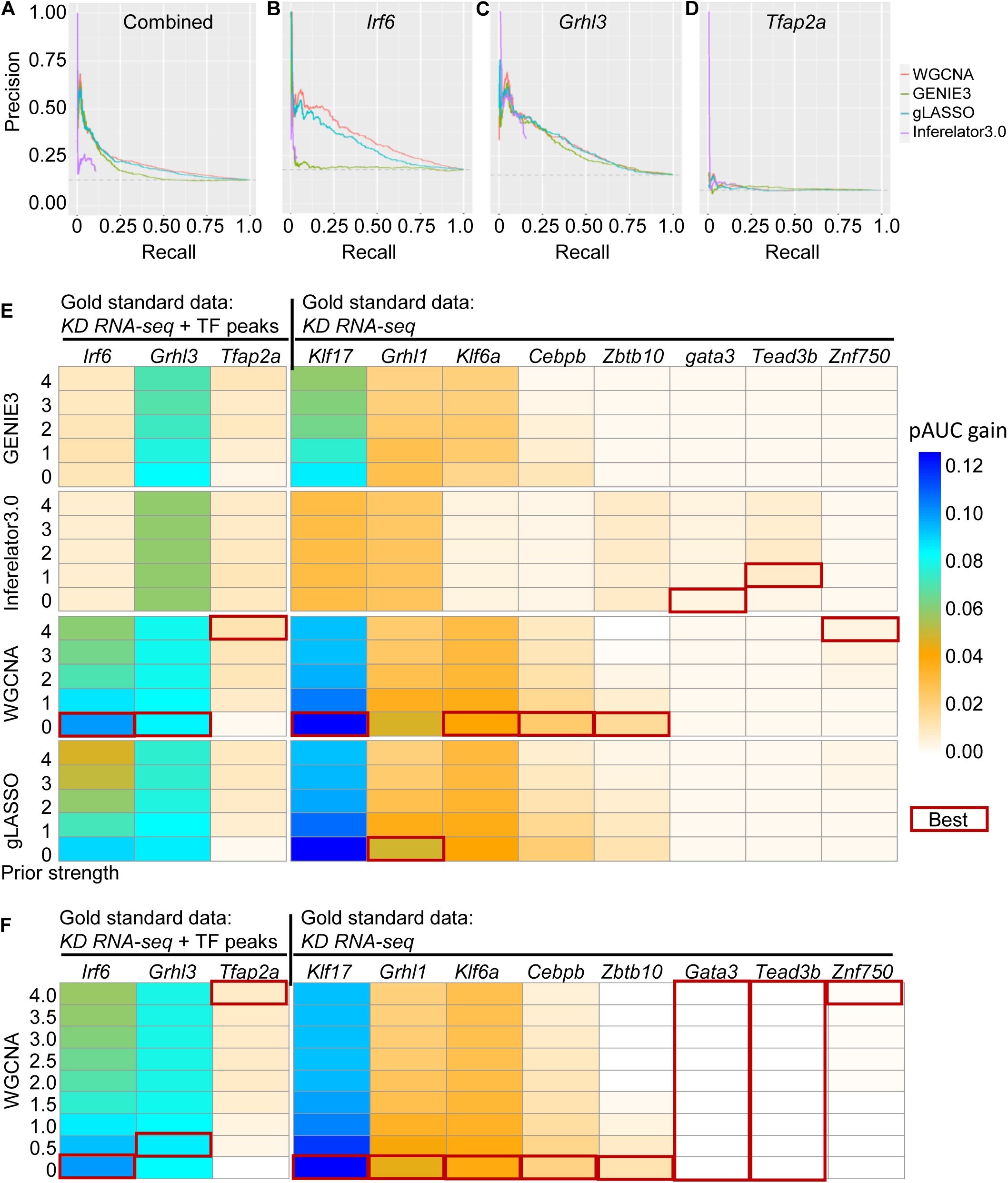
Including ATAC-seq-based prior modestly improves the performance of computational EVL TRN models. Precision recall curves of EVL TRNs generated by four algorithms and with distinct levels of weight given to the prior for A) Combined (all three TFs: Irf6, Grhl3, Tfap2a), B) Irf6, C) Grhl3, and D) Tfap2a. The curves show the ability of the TRNs to recover gold standard edges inferred from ChIP-seq together with RNA-seq of embryos depleted of the TF. The dotted line is performance based on chance. E) Heat maps portraying the gain in partial area under the precision recall curves for each TF and for each algorithm using different values for the weight given to the prior. The optimal performance is on average achieved with a low but non-zero value for the prior weight. F) Heat maps portraying the gain in partial area under the precision recall curves for each TF in WGCNA TRN model.

### Sub-network modules in the EVL TRN

Within the top-performing model, we applied a community detection algorithm to identify three modules of connected genes (**Figure 5A**). Interestingly, each module corresponded to different levels of enrichment of gene expression in EVL cells relative to other cell types (**Figure 5B**). For instance, in module one the average expression of TF-encoding genes was higher in EVL cells than in other cell types (**Figure 5B**). As might be expected, most of the known regulators of periderm differentiation were part of module one (**Table S26**). In summary, subsets of genes in the larger network formed more tightly connected sub-networks or modules, and module one included all the previously described regulators of periderm differentiation.

**Figure 5:**
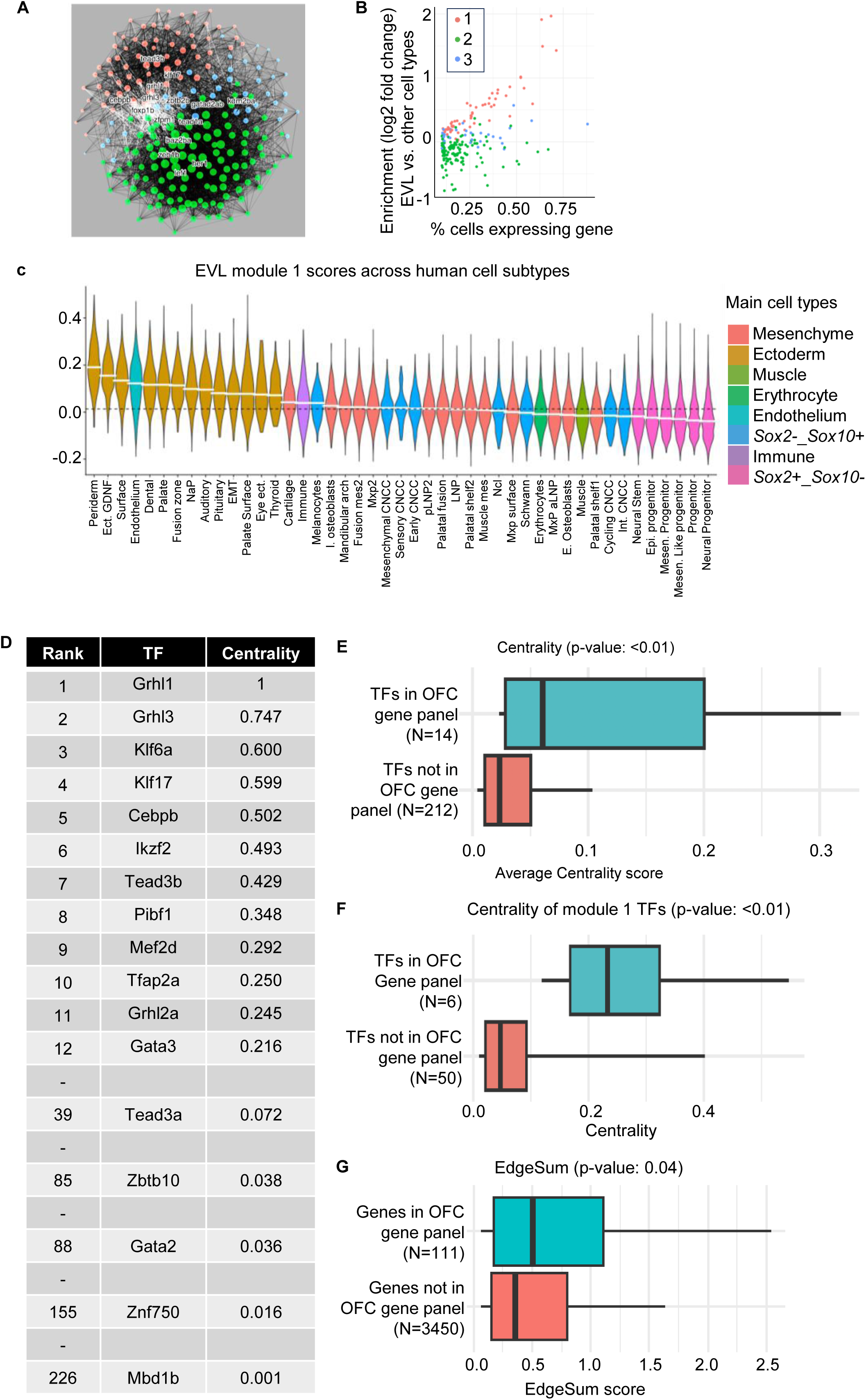
Known EVL-specific TFs are enriched in a single module of best-performing (WGCNA) EVL TRN model. A) Representation of node and edge distribution in three network modules (red, green, and blue). Red module was identified as EVL-specific based on enrichment of known EVL TFs in the module. B) Dot plot showing percent of EVL cells expressing a TF (x-axis) and its expressional enrichment (log2-fold change) in EVL cell cluster versus other cell clusters (y-axis). Color of dots represents the predicted membership of TFs in red, green or blue modules. C) Violin plot of module scores of orthologous genes between human and zebrafish from EVL module 1 across all human craniofacial cell subtypes. D) List of few EVL-specific TFs with their corresponding rank and centrality scores from best-performing WGCNA TRN model. E) Highly connected TFs are significantly enriched for known OFC-associated TFs compared to sparsely connected TFs in the WGCNA based TRN model. F) OFC-associated TFs in module 1 (EVL-specific) are highly connected compared to non-OFC- associated TFs. G) OFC-associated genes are highly connected in EVL TRN model compared to non-OFC-associated TFs.

We next evaluated module preservation of the three EVL modules compared to 45 clusters identified in a single-cell RNA seq analysis of faces explanted from human embryos at Carnegie stage 16 ^77^. As expected, the periderm cluster had the strongest similarity to module 1 in zebrafish EVL TRN (**Figure 5C)**; the periderm cluster and other epithelial clusters were highly similar clusters to zebrafish modules 2 and 3, respectively (**Figure S4A, B**). We recently showed that embryonic facial epithelial clusters were enriched for de novo and rare variants associated with OFC ^77^. We found this enrichment to be higher for the periderm cluster relative to other epithelial clusters (**Fig5C**; **Figure 4A, B**).

### Genes in an OFC risk panel have higher centrality than genes not in the panel

We ranked all the TFs in the target gene list based on their centrality scores. TFs previously reported to regulate periderm differentiation, including Grhl1, Grhl3, Klf17, and Cebpb, were in the top ten most highly ranked TFs (**Figure 5D; Table S26**). Other TFs in the top-ten list, including Tfap2a and Tead3b, were also highly ranked in a recent evaluation of the EVL TRN using single cell multiomic data and the SCENIC+ algorithm ^78^. Irf6, by contrast, although known to be a key regulator of EVL differentiation ^34,35^, was ranked at a middle level, perhaps because high levels of maternally deposited *irf6* mRNA reduces the correlation of *irf6* expression levels with that of other EVL regulators.

We predicted that genes associated with OFC would be among the more highly connected genes in the EVL TRN model. We tested this hypothesis in genes encoding TFs as well as all genes. Based on a review of the literature, we composed a list of 498 zebrafish genes orthologous to genes associated with OFC in mice or humans (**Table S27**). There were 125 OFC-associated genes in our target gene list, of which 14 encoded TFs. The average centrality score (**Table S26;** see Methods section) of OFC-associated TFs was significantly higher than that of non-OFC-associated TF (**Figure 5E**). Among TFs in module one, there was even greater differential in the centrality of OFC-associated vs. non-OFC-associated ones (**Figure 5F**). Because non-TFs have low centrality by definition, we ranked the connectedness of target genes (which can either be TFs or not) by a metric we call *edgeSum* (Methods). Target genes on the list of OFC-associated genes had a modestly higher average edgeSum score than the non-OFC-associated target genes (**Figure 5G**).

We hypothesized that knockdown of highly connected TFs would recapitulate the phenotype of either *irf6* or *grhl3* mutants. When synchronized wild-type embryos reach 6 hpf (**Figure 6A**), maternal *irf6* mutant embryos derived from a mother homozygous for a strong loss-of-function allele (and wild-type embryos injected with mRNA encoding a constitutive repressor variant of Irf6) remain arrested at 4 hpf (sphere stage) and rupture while wild-types reach 8 hpf (late gastrula stage) (**Figure 6B**); EVL differentiation appears to fail utterly in such embryos ^34,35^. Interestingly, zygotic *irf6* mutants and *irf6* morphants are viable implying that maternal Irf6 suffices for its requirement in EVL development ^34,35^. The phenotype of embryos homozygous for loss-of-function alleles of *grhl3* depends on as-of-yet undetermined variable quality of the housing conditions; in certain conditions they die at about 11 hpf (mid-somitogenesis) with rupture of the periderm ^36,79^. In other conditions *grhl3* zygotic mutants live until about 10 days with variably penetrant phenotypes ^37^. Importantly, in our hands *grhl3* morphants do not rupture ^80^, probably because the morpholino does not completely eliminate *grhl3* expression; by contrast, about 25% of *grhl3* crispants die at about 11 hpf (**Figure S5A, B**). We injected morpholinos (MOs) targeting genes encoding seven top-ranked TFs (Grhl1, Grhl3, Klf17, Cebpb, Tead3b, Klf6a, and Tfap2a), two middle-ranked TFs (Zbtb10 and Gata3), and a lowly ranked one (Znf750); all of these TFs are in module one. We evaluated injected embryos up to 24 hpf. In no case did morphant embryos rupture.

**Figure 6:**
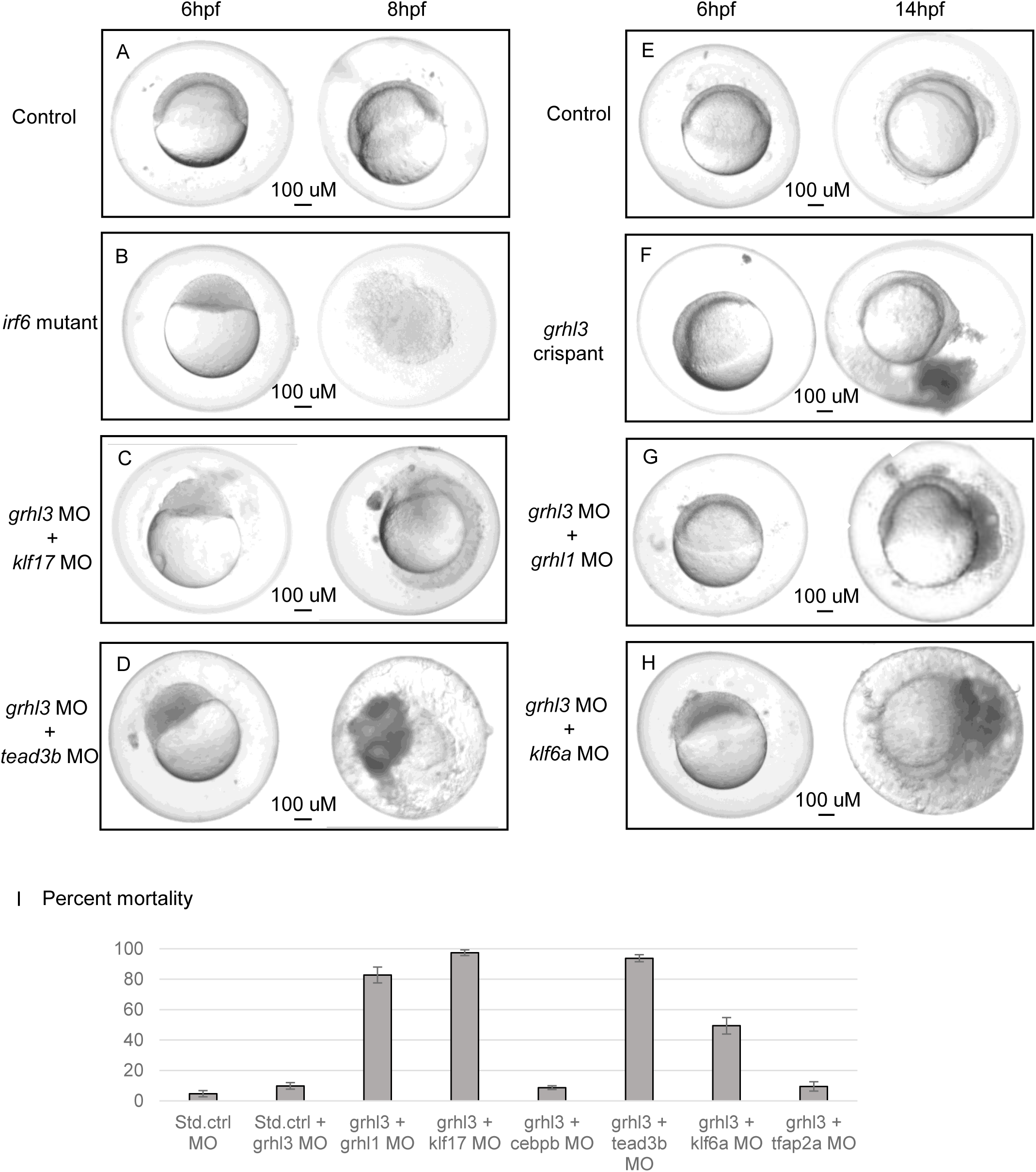
Experimental functional validation of highly connected TFs identified in best performing WGCNA-based TRN model. A-H) Phenotyping of zebrafish morphant embryos. Representative images of zebrafish embryos injected with standard control and/ or combination of gene-specific morpholinos. I) Percent mortality observed in zebrafish embryos injected with standard control and/ or combination of gene-specific morpholinos.

Next, because two studies found that embryos injected with *grhl3* MO were sensitized to other manipulations affecting EVL or skin development ^80,81^, we co-injected the *grhl3* MO with MO targeting the TFs mentioned above and observed four distinct phenotypes. First, embryos injected with the combination of *grhl3* MO and either the *klf17* MO (**Figure 6C**) or *tead3b* MO (**Figure 6D**) arrested at 4 hpf and ruptured at about 8 hpf, closely mimicking the phenotype of *irf6* maternal mutants. Second, embryos injected with the *grhl3* MO and the *grhl1* MO ruptured during somitogenesis (**Figure 6G**) similar to *grhl3* mutants ^36,79^, as we have reported before ^80^. Third, the combination of *grhl3* MO and *klf6a* MO yielded a phenotype of cell extrusion by 8 hpf (**Figure 6H**). Finally, co-injection of *grhl3* MO with high-ranking TFs *cebpb* or *tfap2a* did not yield an overt phenotype. Similarly, co-injection of *grhl3* MO with those targeting gene encoding middle-ranking TFs (Zbtb10 and Gata3) or low-ranking TFs (Znf750), did not visibly affect periderm or skin. Co-injection of *grhl3* Morpholino with those targeting *grhl1*, *klf17*, *tead3b*, or *klf6a* significantly increased the frequency of periderm rupture phenotype in injected embryos (**Figure 6I**).

To confirm the MO induced phenotypes, we next injected wild-type embryos with Cas9 protein and guide RNAs targeting the highly ranked TFs, alone or with gRNAs targeting *grhl3* and evaluated injected embryos up until 24 hpf (**Figure S5A-E**). While approximately 25 percent of embryos injected with gRNAs targeting *grhl3* and Cas9 protein (i.e., *grhl3* crispants) ruptured prior to 24 hpf, the other TF crispants had mortality comparable to control crispants (i.e., injected with a non-targeting gRNA and Cas9 protein) and did not exhibit a grossly observable periderm phenotype. However, combining guide RNAs targeting *grhl3* with those targeting *grhl1*, *klf17*, *tead3b*, or *klf6a* significantly increased the frequency of periderm rupture phenotype in injected embryos (**Figure S5F**). Overall, there was a significant association between the centrality of a TF in the network and phenotypic consequences (p=0.02, Wilcoxon rank sum test), with higher ranking TFs more likely to produce an overt phenotype upon being knocked down in combination with *grhl3*.

### Rare variants in GRHL1 are present in individuals with OFC

We examined sequence data for 798 cleft lip (CL), 1062 cleft lip and palate (CLP), and 554 cleft palate (CP) cases (primarily trios) for rare variants of interest, defined as having an allele frequency less than 0.1% in any gnomAD population, and being predicted to alter protein function (e.g., missense, inframe indels, nonsense, frameshift, splice), in the principal transcript of *GRHL1*, *KLF6*, *KLF17*, and *TEAD3*. Interestingly, in *GRHL1* we identified 24 total variants fulfilling those criteria (**Figure S6**). Of these, 15 had a CADD score greater than 25 (**Figure 7A**), indicating they are predicted to be in the top 0.3% of the most deleterious substitutions ^82^. We found one *de novo* variant, one inherited from an affected mother, and the rest (i.e., 13 variants) inherited from unaffected parents **(Table S28**). Of these, we considered five variants to be very likely to cause deleterious effects because they were nonsense variants either absent from the gnomAD database or ultrarare (observed only 4 times in gnomAD) or, for one missense variant, transmitted from an affected parent (**Figure 7B**).

**Figure 7:**
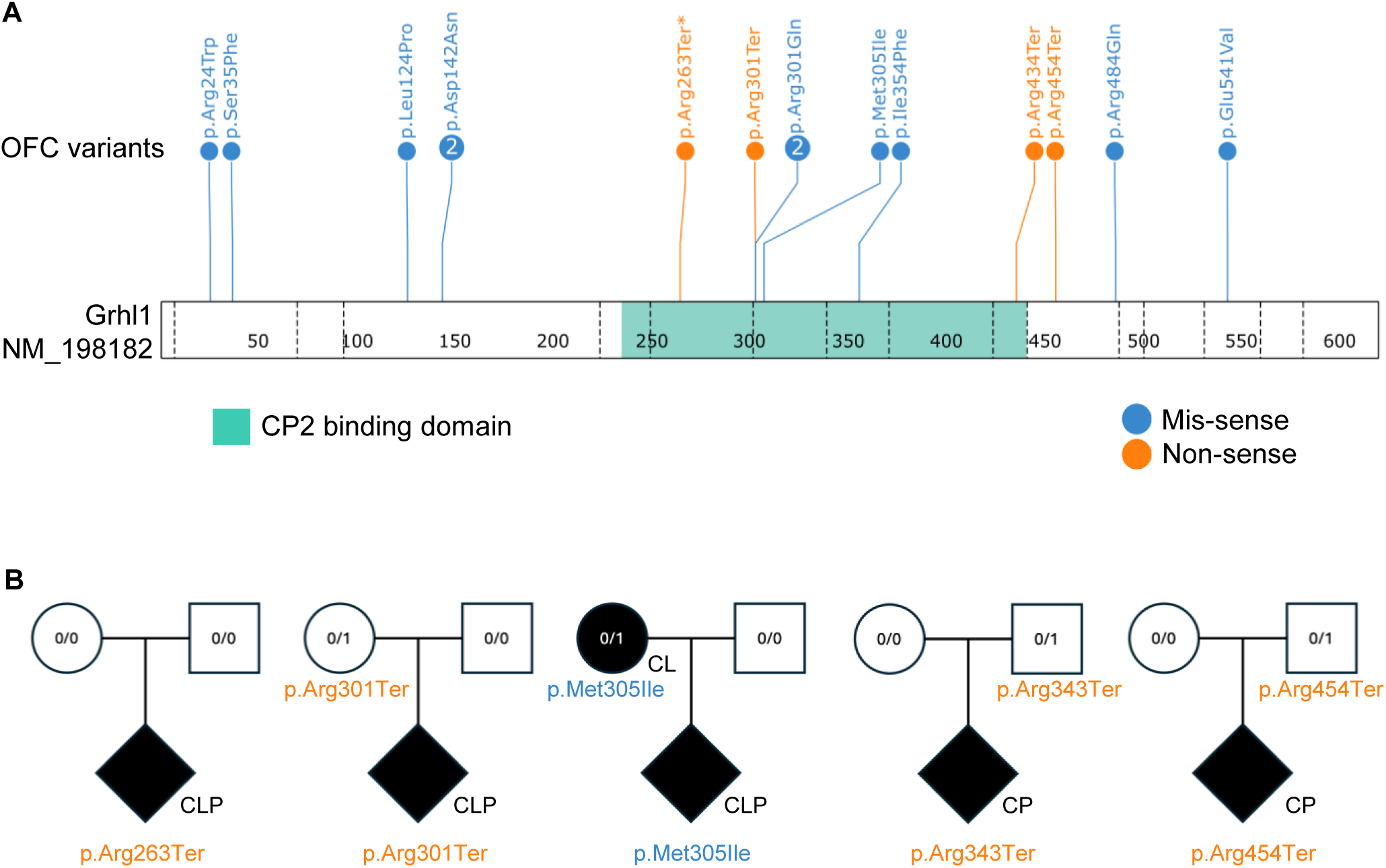
Rare protein-altering variants with prediction of deleterious effects in *GRHL1*. A) Lollipop plot of 15 variants with minor allele frequency <0.1% in gnomAD v4.1 and CADD score >25. The asterisk indicates a *de novo* variant. The black arrows denote variants with high likelihood of deleterious effects and lower-case letters indicate inheritance patterns as shown in panel B. B) Pedigree depiction for each predicted deleterious variant shown in panel A.

When considering all *GRHL1* variants (MAF <0.1% and protein-altering), the difference in transmission rate was not significant at p=0.215 (transmitted n=20, untransmitted n=12). Nor was it significant when considering variants with CADD > 25 (transmitted n =12, untransmitted n=6; p=0.238). However, if we only consider variants absent in gnomAD, there was a significant difference at p=0.013 (transmitted n=12, untransmitted n=2), suggesting there is an overall burden of ultra-rare variants that extends to other missense variants in *GRHL1*, but larger sample sizes and or functional studies are needed to confirm.

Although we identified rare, protein-altering variants in *KLF6*, *KLF17*, and *TEAD3*, none reached the same evidence for pathogenicity as the *GRHL1* variants **(Table S28**). Finally, because variants in genes of this TRN are associated with Van der Woude syndrome, we asked whether variants in *GRHL1*, *KLF6*, *KLF17*, and *TEAD3* could explain any of the unsolved individuals who lack causal variants in the known genes (*IRF6*, *GRHL3*, and *PRKCI*) ^26,27,29^. We sequenced 12 such individuals and found no variants of interest.

## Discussion

In this study, we employed a multi-modal systems biology approach to bridge the gap between periderm development and the genetic architecture of OFC condition. By integrating single-cell multiome (ATAC + RNA-seq) with computational inference of network models, we identified a core transcriptional regulatory network (TRN) governing periderm differentiation in zebrafish. A key strength of this study was the functional validation of network hubs. While single-gene knockdowns of high-centrality TFs did not result in overt phenotypes, sensitized screening in embryos depleted of *grhl3* revealed genetic synergies. For instance, simultaneous reduction of *grhl3* with *tead3b* or with *klf17* mimicked the severe *irf6* maternal mutant phenotype suggesting these factors cooperate within a tightly regulated hierarchy to ensure periderm integrity. Klf17 has previously been implicated in zebrafish periderm differentiation ^83,84^, and Tead3 binding sites are enriched in periderm enhancers ^73^. Simultaneous reduction of *grhl3* and of *grhl1* recapitulated the phenotype of *grhl3* loss of function mutants, and simultaneous reduction of *grhl3* and *klf6a* led to blebbing and rupture. These functional tests confirm the ability of TRN analysis to discover novel regulators of cellular differentiation.

A second strength of the study was the use of gold-standard datasets to benchmark network performance, which showed that incorporating chromatin accessibility as prior evidence yielded only a modest gain in network performance over models based solely on correlated gene expression. In a previous study also using gold-standard datasets ^44^, a position weight matrix (PWM)-based prior yielded a stronger improvement in network performance. The lesser benefit of prior in our study may reflect, first, that ATAC-seq data from single cells are inherently sparse, and the number of periderm cells we evaluated (i.e., several hundred) may have been insufficient to reveal all the periderm cis-regulatory elements; we believe our efforts to use SCENIC+ were foiled for this reason ^85^. Second, PWMs for TF binding sites were available for only 151 of the 226 TFs expressed in EVL, whereas the study mentioned above was conducted in human cells where the catalog of PWMs is more complete. Third, a given PWM can correspond to more than one TF, lessening the value of the prior matrix for inferring edges. This issue may have a bigger effect in teleost fish than in mammals because of a genome duplication event in the former ^86^. Finally, PWMs are a poor proxy for TF peaks, because they are present in the genome in higher numbers than corresponding ChIP-seq peaks and, conversely, because they are not present beneath every ChIP-seq peak ^87^. New machine learning tools can infer TF binding events from DNA sequence with better accuracy than PWMs ^88^; these tools may improve the value of incorporating prior data in TRN modeling. Despite the limitations of PWM-based prior, a recent study utilizing machine learning to infer PWMs for zebrafish TFs and relying heavily on prior evidence accurately captured the importance of Irf6 in the EVL TRN ^78^, while our EVL TRN models, which relied primarily on correlated-gene-expression evidence, did not. The two studies otherwise largely agree on the list of key transcriptional regulators of EVL, although the prior-based study to not identify Grhl1 to be among them, which our functional tests clearly establish it to be.

Unexpectedly, the pair-wise evaluation of correlated gene expression used in WGCNA and in gLASSO with alpha set to zero yielded better-performing network models than the more complex regression modes used in the other algorithms ^44,50,52,89^. It was recently reported that the EVL TRN is relatively shallow, with a small number of key regulators governing expression of a large number of effector genes, relative to TRNs regulating later developing cell types, which have more layers of transcription factors ^78^. It is possible that WGCNA and similar tools work best on shallow networks while tools using regression will perform better on deeper ones. Testing this possibility will require relevant gold-standard datasets, which may be more challenging for later-developing cell types than in it was for the EVL, where we could simply knockdown a transcription factor in the whole embryo and harvest RNA when EVL was virtually the only cell type present.

Turning to potential connections to human genetics, we found that the zebrafish EVL most closely resembles the periderm among all cell subtypes in a single cell sequencing analysis of human embryonic faces ^77^, revealing a conserved circuitry for this essential barrier tissue. Emphasizing the clinical importance of this circuitry, among TFs whose orthologs are expressed in zebrafish EVL, those on a panel of OFC-associated genes had a higher average centrality score in the EVL TRN than those not on the panel. For instance, *GRHL3*, a known contributor to VWS ^27^ and nsOFC ^8,90^, was second-highest ranking. In addition, we found that periderm is enriched, relative to other epithelial clusters, for de novo variants in OFC patients. Finally, we examined the *GHRL1, KLF6, KLF17* and *TEAD3* genes in whole genome sequence data from OFC trios and found that ultrarare mutations in *GRHL1* are over-transmitted to cases. There is now evidence of a burden of variants in nsOFC patients, or in VWS patients, in orthologs of four key regulators of zebrafish EVL differentiation (*GRHL1, GRHL3, IRF6* and *PRKCI*) ^8,26,27,29,90,91^; however we did not find evidence for such a burden in three others (*KLF6*, *KLF17*, or *TEAD3).* It is possible that mutations in these genes result in one of hundreds of other syndromes that include OFC (http://www.ncbi.nlm.nih.gov/OMIM), and variants near these gene may interact with those near *GRHL3* to make the phenotype of peridermopathies more severe. All are expressed in human oral periderm at Carnegie Stage 20 ^77^, which is a few days before palate shelves fuse. KLF17 interacts with TGFβ/SMAD signaling and p53 to suppress tumor progression ^92,93^, and inhibits epithelial-to-mesenchymal transition ^94,95^. It also regulates pluripotency ^96^. TEAD3 cooperates with TEAD1 to promote epidermal differentiation ^97^, and to drive human embryonic stem cells to become surface ectoderm in vitro ^98^. An individual in the DECIPHER database has a de novo missense variant in *TEAD3* with unspecified abnormalities of the head and neck, digestive system, musculoskeletal system, and nervous system ^99^. Whether or not orthologs of all key regulators of EVL differentiation are OFC risk genes, this study emphasizes a shared TRN between zebrafish EVL and human oral periderm and that disruption of this TRN underlies many cases of orofacial cleft.

## Acknowledgements

The authors would like to thank Greg Bonde for helping with morpholino injections at an early phase of this project and fish husbandry at the University of Iowa, and Frank Radella for running the zebrafish facility at the University of Washington.

## Author contributions

Conceptualization: RAC, SKS, PB, EJLC; Methodology: SKS, EJLC, CK, LP, KMD, EBL; Formal analysis: SKS, PB, AH, SWC, KR, AM, JC; Investigation: Writing - original draft: SKS; Writing - review & editing: RAC, SKS, PB, EJLC.; Visualization: SKS, PB, AH; Supervision: RAC; Funding acquisition: RAC, EJLC, EL.

## Data availability

The sequencing data generated in this study were submitted to NCBI GEO (https://www.ncbi.nlm.nih.gov/geo/). Whole genome sequencing datasets are available from the Kids First Data Resource Center and dbGaP (accession numbers: phs002220.v1.p1, phs001168.v1.p1, phs001420.v1.p1, phs001997.v1.p1, phs002595.v1.p1).

## Supplemental information

**Figure S1**: Single cell multiome (ATAC + RNA-seq) on zebrafish shield-stage embryos. UMAP plots based on A) scRNA-seq, B) scATAC-seq, and C) combined (scRNA + scATAC) data. D) Bubble plot showing top 5 marker genes from each cell cluster and their corresponding expression.

**Figure S2**: Genome viewer tracks of RNA-seq data from morpholino-injected zebrafish embryos showing splice-blocking in corresponding transcripts. A) *gata3* MO. B) *grhl3* MO. C) *tfap2a* MO. D) *tead3b* MO. C) *zbtb10* MO.

**Figure S3**: Genomic annotations of called TF-binding peaks. A) Irf6 ChIP-seq peaks. B) Grhl3 CUT&RUN peaks. C) Tfap2a CUT&RUN peaks.

**Figure S4**: Enrichment of zebrafish EVL module genes in human relevant subtypes and association with OFC risk. A) EVL module 2. B) EVL module 3.

**Figure S5**: Phenotyping of zebrafish crispant embryos. A-E’) Representative images of zebrafish embryos injected with CRISPR-Cas9 guides (scale bar = 100 uM). F) Percent mortality observed in zebrafish embryos injected with CRISPR-Cas9 guides.

**Figure S6**: All rare protein-altering variants with prediction of deleterious effects in *GRHL1*.

## References

1. Dixon, M.J., Marazita, M.L., Beaty, T.H., and Murray, J.C. (2011). Cleft lip and palate: understanding genetic and environmental influences. Nat Rev Genet 12, 167–178. 10.1038/nrg2933.

2. Bryan, E., Little, J., and Burn, J. (1987). 10 Congenital Anomalies in Twins. Baillière’s Clinical Obstetrics and Gynaecology, Fetal Diagnosis of Genetic Defects 1, 697–721. 10.1016/S0950-3552(87)80012-3.

3. Grosen, D., Bille, C., Petersen, I., Skytthe, A., Hjelmborg, J.v.B., Pedersen, J.K., Murray, J.C., and Christensen, K. (2011). Risk of Oral Clefts in Twins. Epidemiology 22, 313–319. 10.1097/EDE.0b013e3182125f9c.

4. Beaty, T.H., Murray, J.C., Marazita, M.L., Munger, R.G., Ruczinski, I., Hetmanski, J.B., Liang, K.Y., Wu, T., Murray, T., Fallin, M.D., et al. (2010). A genome-wide association study of cleft lip with and without cleft palate identifies risk variants near MAFB and ABCA4. Nat Genet 42, 525–529. 10.1038/ng.580.

5. Bishop, M.R., Perez, K.K.D., Sun, M., Ho, S., Chopra, P., Mukhopadhyay, N., Hetmanski, J.B., Taub, M.A., Moreno-Uribe, L.M., Valencia-Ramirez, L.C., et al. (2020). Genome-wide Enrichment of De Novo Coding Mutations in Orofacial Cleft Trios. The American Journal of Human Genetics 107, 124–136. 10.1016/j.ajhg.2020.05.018.

6. He, M., Zuo, X., Liu, H., Wang, W., Zhang, Y., Fu, Y., Zhen, Q., Yu, Y., Pan, Y., Qin, C., et al. (2020). Genome-wide Analyses Identify a Novel Risk Locus for Nonsyndromic Cleft Palate. J Dent Res 99, 1461–1468. 10.1177/0022034520943867.

7. Leslie, E.J., Carlson, J.C., Shaffer, J.R., Feingold, E., Wehby, G., Laurie, C.A., Jain, D., Laurie, C.C., Doheny, K.F., McHenry, T., et al. (2016). A multi-ethnic genome-wide association study identifies novel loci for non-syndromic cleft lip with or without cleft palate on 2p24.2, 17q23 and 19q13. Hum Mol Genet 25, 2862–2872. 10.1093/hmg/ddw104.

8. Leslie, E.J., Liu, H., Carlson, J.C., Shaffer, J.R., Feingold, E., Wehby, G., Laurie, C.A., Jain, D., Laurie, C.C., Doheny, K.F., et al. (2016). A Genome-wide Association Study of Nonsyndromic Cleft Palate Identifies an Etiologic Missense Variant in GRHL3. The American Journal of Human Genetics 98, 744–754. 10.1016/j.ajhg.2016.02.014.

9. 9. Mangold, E., Ludwig, K.U., Birnbaum, S., Baluardo, C., Ferrian, M., Herms, S., Reutter, H., de Assis, N.A., Chawa, T.A., Mattheisen, M., et al. (2010). Genome-wide association study identifies two susceptibility loci for nonsyndromic cleft lip with or without cleft palate. Nat Genet 42, 24–26. 10.1038/ng.506.

10. Mukhopadhyay, N., Feingold, E., and Moreno-Uribe, L. (2021). Genome-Wide Association Study of Non-Syndromic Orofacial Clefts in a Multiethnic Sample of Families and Controls Identifies Novel Regions. Front Cell Dev Biol 9. 10.3389/fcell.2021.621482.

11. Génin, E. (2020). Missing Heritability of Complex Diseases: Case Solved? Hum Genet 139, 103–113. 10.1007/s00439-019-02034-4.

12. Tabangin, M.E., Woo, J.G., and Martin, L.J. (2009). The Effect of Minor Allele Frequency on the Likelihood of Obtaining False Positives. BMC Proceedings 3 *Suppl 7*. 10.1186/1753-6561-3-s7-s41.

13. Beaty, T.H., Marazita, M.L., and Leslie, E.J. (2016). Genetic Factors Influencing Risk to Orofacial Clefts: Today’s Challenges and Tomorrow’s Opportunities. F1000Research 5. 10.12688/f1000research.9503.1.

14. Rahimov, F., Jugessur, A., and Murray, J.C. (2012). Genetics of Nonsyndromic Orofacial Clefts. The Cleft Palate-Craniofacial Journal : Official Publication of the American Cleft Palate-Craniofacial Association 49, 73–91. 10.1597/10-178.

15. Koboldt, D.C., Ding, L., Mardis, E.R., and Wilson, R.K. (2010). Challenges of Sequencing Human Genomes. Briefings in Bioinformatics 11, 484–498. 10.1093/bib/bbq016.

16. Musunuru, K., Pirruccello, J.P., and Do, R. (2010). Exome Sequencing, ANGPTL3 Mutations, and Familial Combined Hypolipidemia. N Engl J Med 363, 2220–2227. 10.1056/NEJMoa1002926.

17. Ng, S.B., Turner, E.H., and Robertson, P.D. (2009). Targeted Capture and Massively Parallel Sequencing of 12 Human Exomes. Nature 461. 10.1038/nature08250.

18. Marsh, J.A., and Teichmann, S.A. (2023). Predicting Pathogenic Protein Variants. Science 381, 6664. 10.1126/science.adj8672.

19. Hammond, N.L., and Dixon, M.J. (2022). Revisiting the embryogenesis of lip and palate development. Oral Diseases 28, 1306–1326. 10.1111/odi.14174.

20. Richardson, R.J., Hammond, N.L., Coulombe, P.A., Saloranta, C., Nousiainen, H.O., Salonen, R., Berry, A., Hanley, N., Headon, D., Karikoski, R., and Dixon, M.J. (2014). Periderm prevents pathological epithelial adhesions during embryogenesis. J. Clin. Invest. 124, 3891–3900. 10.1172/JCI71946.

21. Yoshida, M., Shimono, Y., and Togashi, H. (2012). Periderm Cells Covering Palatal Shelves Have Tight Junctions and Their Desquamation Reduces the Polarity of Palatal Shelf Epithelial Cells in Palatogenesis. Genes to Cells 17, 455–472. 10.1111/j.1365-2443.2012.01601.x.

22. Kashgari, G., Meinecke, L., Gordon, W., Ruiz, B., Yang, J., Ma, A.L., Xie, Y., Ho, H., Plikus, M.V., Nie, Q., et al. (2020). Epithelial Migration and Non-adhesive Periderm Are Required for Digit Separation during Mammalian Development. Developmental Cell 52, 764–778.e764. 10.1016/j.devcel.2020.01.032.

23. Findlater, G.S., McDougall, R.D., and Kaufman, M.H. (1993). Eyelid development, fusion and subsequent reopening in the mouse. J Anat 183, 121–129.

24. Maconnachie, E. (1979). A Study of Digit Fusion in the Mouse Embryo. Development 49, 259–276. 10.1242/dev.49.1.259.

25. 25. Khandelwal, K.D., van den Boogaard, M.-J.H., Mehrem, S.L., Gebel, J., Fagerberg, C., van Beusekom, E., van Binsbergen, E., Topaloglu, O., Steehouwer, M., Gilissen, C., et al. (2019). Deletions and loss-of-function variants in TP63 associated with orofacial clefting. Eur J Hum Genet 27, 1101–1112. 10.1038/s41431-019-0370-0.

26. 26. Kondo, S., Schutte, B.C., Richardson, R.J., Bjork, B.C., Knight, A.S., Watanabe, Y., Howard, E., Ferreira de Lima, R.L.L., Daack-Hirsch, S., Sander, A., et al. (2002). Mutations in IRF6 cause Van der Woude and popliteal pterygium syndromes. Nat Genet 32, 285–289. 10.1038/ng985.

27. Peyrard-Janvid, M., Leslie, E.J., Kousa, Y.A., Smith, T.L., Dunnwald, M., Magnusson, M., Lentz, B.A., Unneberg, P., Fransson, I., Koillinen, H.K., et al. (2014). Dominant mutations in GRHL3 cause Van der Woude Syndrome and disrupt oral periderm development. Am J Hum Genet 94, 23–32. 10.1016/j.ajhg.2013.11.009.

28. Chalmers, A.D., Strauss, B., and Papalopulu, N. (2003). Oriented cell divisions asymmetrically segregate aPKC and generate cell fate diversity in the early Xenopus embryo. Development 130, 2657–2668. 10.1242/dev.00490.

29. Robinson, K., Singh, S.K., Walkup, R.B., Fawwal, D.V., Vilfort, K.M., Koloskee, A., Fashina, A., Adeyemo, W.L., Beaty, T.H., Butali, A., et al. (2025). Rare variants in PRKCI cause Van der Woude syndrome and other features of peridermopathy. Am J Hum Genet 112, 2422–2439. 10.1016/j.ajhg.2025.08.008.

30. Chang, W.-J., and Hwang, P.-P. (2011). Development of Zebrafish Epidermis. Birth Defects Research Part C: Embryo Today: Reviews 93, 205–214. 10.1002/bdrc.20215.

31. Kimmel, C.B., Ballard, W.W., Kimmel, S.R., Ullmann, B., and Schilling, T.F. (1995). Stages of Embryonic Development of the Zebrafish. Developmental Dynamics 203, 253–310. 10.1002/aja.1002030302.

32. Warga, R.M., and Kimmel, C.B. (1990). Cell Movements during Epiboly and Gastrulation in Zebrafish. Development 108, 569–580. 10.1242/dev.108.4.569.

33. 33. Carroll, S.H., Trevino, C.M., and Li, E.B. (2020). An Irf6-Esrp1/2 Regulatory Axis Controls Midface Morphogenesis in Vertebrates.

34. Li, E.B., Truong, D., Hallett, S.A., Mukherjee, K., Schutte, B.C., and Liao, E.C. (2017). Rapid functional analysis of computationally complex rare human IRF6 gene variants using a novel zebrafish model. PLOS Genetics 13, e1007009. 10.1371/journal.pgen.1007009.

35. 35. Sabel, J.L., d’Alençon, C., O’Brien, E.K., Van Otterloo, E., Lutz, K., Cuykendall, T.N., Schutte, B.C., Houston, D.W., and Cornell, R.A. (2009). Maternal Interferon Regulatory Factor 6 is required for the differentiation of primary superficial epithelia in Danio and Xenopus embryos. Developmental Biology 325, 249–262. 10.1016/j.ydbio.2008.10.031.

36. Miles, L.B., Darido, C., Kaslin, J., Heath, J.K., Jane, S.M., and Dworkin, S. (2017). Mis-Expression of Grainyhead-like Transcription Factors in Zebrafish Leads to Defects in Enveloping Layer (EVL) Integrity, Cellular Morphogenesis and Axial Extension. Scientific Reports 7, 1. 10.1038/s41598-017-17898-7.

37. 37. Mathiyalagan, N., Johnson, T.K., Di Pastena, Z., Fuller, J.N., Miles, L.B., and Dworkin, S. (2025). Loss of the epithelial transcription factor grhl3 leads to variably penetrant developmental phenotypes in zebrafish. Dev Dyn 254, 1133–1147. 10.1002/dvdy.70003.

38. Fukazawa, C., Santiago, C., and Park, K.M. (2010). Poky/Chuk/Ikk1 Is Required for Differentiation of the Zebrafish Embryonic Epidermis. Developmental Biology 346, 2. 10.1016/j.ydbio.2010.07.037.

39. Davidson, E., and Levin, M. (2005). Gene Regulatory Networks. Proceedings of the National Academy of Sciences of the United States of America 102, 4935. 10.1073/pnas.0502024102.

40. Davidson, E.H., McClay, D.R., and Hood, L. (2003). Regulatory Gene Networks and the Properties of the Developmental Process. Proceedings of the National Academy of Sciences 100, 4. 10.1073/pnas.0437746100.

41. Howard, M.L., and Davidson, E.H. (2004). Cis-Regulatory Control Circuits in Development. Developmental Biology 271, 1. 10.1016/j.ydbio.2004.03.031.

42. Huynh-Thu, V.A., and Sanguinetti, G. (2019). Gene Regulatory Network Inference: An Introductory Survey. In Gene Regulatory Networks: Methods and Protocols, G. Sanguinetti, ed. (and Vân Anh Huynh-Thu. Springer).

43. Saint-Antoine, M.M., and Singh, A. (2020). Network Inference in Systems Biology: Recent Developments, Challenges, and Applications. Current Opinion in Biotechnology, Nanobiotechnology ● Systems Biology 63. 10.1016/j.copbio.2019.12.002.

44. 44. Miraldi, E.R., Pokrovskii, M., Watters, A., Castro, D.M., De Veaux, N., Hall, J.A., Lee, J.Y., Ciofani, M., Madar, A., Carriero, N., et al. (2019). Leveraging chromatin accessibility for transcriptional regulatory network inference in T Helper 17 Cells. Genome Res 29, 449–463. 10.1101/gr.238253.118.

45. Badia, I.M.P., Wessels, L., Muller-Dott, S., Trimbour, R., Ramirez Flores, R.O., Argelaguet, R., and Saez-Rodriguez, J. (2023). Gene regulatory network inference in the era of single-cell multi-omics. Nat Rev Genet 24, 739–754. 10.1038/s41576-023-00618-5.

46. Hao, Y., Hao, S., and Andersen-Nissen, E. (2021). Integrated Analysis of Multimodal Single-Cell Data. Cell 184, 3573–3587 3529. 10.1016/j.cell.2021.04.048.

47. Zhang, Y., Liu, T., and Meyer, C.A. (2008). Model-Based Analysis of ChIP-Seq (MACS. Genome Biology 9. 10.1186/gb-2008-9-9-r137.

48. Grant, C.E., Bailey, T.L., and Noble, W.S. (2011). FIMO: Scanning for Occurrences of a given Motif. Bioinformatics. 10.1093/bioinformatics/btr064.

49. Langfelder, P., and Horvath, S. (2008). WGCNA: An R Package for Weighted Correlation Network Analysis. BMC Bioinformatics 9, 559. 10.1186/1471-2105-9-559.

50. Huynh-Thu, V.A., Irrthum, A., Wehenkel, L., and Geurts, P. (2010). Inferring Regulatory Networks from Expression Data Using Tree-Based Methods. PLOS ONE 5. 10.1371/journal.pone.0012776.

51. Friedman, J., Hastie, T., and Tibshirani, R. (2008). Sparse Inverse Covariance Estimation with the Graphical Lasso. Biostatistics 9, 432–441. 10.1093/biostatistics/kxm045.

52. Gibbs, S., Claudia, C.A.J., and Saldi, G.-A. (2022). High-Performance Single-Cell Gene Regulatory Network Inference at Scale: The Inferelator 3.0. Bioinformatics 38, 2519–2528. 10.1093/bioinformatics/btac117.

53. Hoshijima, K., Jurynec, M.J., Shaw, D.K., Jacobi, A.M., Behlke, M.A., and Grunwald, D.J. (2019). Highly Efficient CRISPR-Cas9-Based Methods for Generating Deletion Mutations and F0 Embryos That Lack Gene Function in Zebrafish. Developmental Cell 51, 645–657 644. 10.1016/j.devcel.2019.10.004.

54. Dobin, A., Davis, C.A., and Schlesinger, F. (2013). STAR: Ultrafast Universal RNA-Seq Aligner. Bioinformatics (Oxford, England 29, 15–21. 10.1093/bioinformatics/bts635.

55. Liao, Y., Smyth, G.K., and Shi, W. (2014). featureCounts: an efficient general purpose program for assigning sequence reads to genomic features. Bioinformatics 30, 923–930. 10.1093/bioinformatics/btt656.

56. Love, M.I., Huber, W., and Anders, S. (2014). Moderated estimation of fold change and dispersion for RNA-seq data with DESeq2. Genome Biology 15, 550. 10.1186/s13059-014-0550-8.

57. Stephens, M. (2017). False Discovery Rates: A New Deal. Biostatistics 18, 275–294. 10.1093/biostatistics/kxw041.

58. Li, E.B.H. (2018). Investigating the Roles of IRF6 in Epithelial Maturation, Craniofacial Development, and Orofacial Cleft Pathogenesis.

59. Langmead, B., and Salzberg, S.L. (2012). Fast Gapped-Read Alignment with Bowtie 2. Nature Methods 9, 357–359. 10.1038/nmeth.1923.

60. Pons, P., and Latapy, M. (2005). Computing Communities in Large Networks Using Random Walks. In Computer and Information Sciences - ISCIS 2005, Yolum, and T. Güngör, eds. (Springer).

61. Ware, J.S., Samocha, K.E., Homsy, J., and Daly, M.J. (2015). Interpreting de novo Variation in Human Disease Using denovolyzeR. Curr Protoc Hum Genet 87, 7 25 21-27 25 15. 10.1002/0471142905.hg0725s87.

62. Samocha, K.E., Robinson, E.B., Sanders, S.J., Stevens, C., Sabo, A., McGrath, L.M., Kosmicki, J.A., Rehnstrom, K., Mallick, S., Kirby, A., et al. (2014). A framework for the interpretation of de novo mutation in human disease. Nat Genet 46, 944–950. 10.1038/ng.3050.

63. Bishop, M.R., Perez, K.K.D., and Sun, M. (2020). Genome-Wide Enrichment of De Novo Coding Mutations in Orofacial Cleft Trios. The American Journal of Human Genetics 107, 124–136. 10.1016/j.ajhg.2020.05.018.

64. 64. Curtis, S.W., Yang, C., Sanchis-Juan, A., Singleton, K., Beaty, T.H., Erger, F., Epstein, M.P., Feingold, E., Krause, M., Uribe, L.M.M., et al. (2025). Haploinsufficiency of GRHL2 is associated with orofacial clefting in humans.

65. Robinson, K., Mosley, T.J., Rivera-Gonzalez, K.S., Jabbarpour, C.R., Curtis, S.W., Adeyemo, W.L., Beaty, T.H., Butali, A., Buxo, C.J., Cutler, D.J., et al. (2023). Trio-based GWAS identifies novel associations and subtype-specific risk factors for cleft palate. HGG Adv 4, 100234. 10.1016/j.xhgg.2023.100234.

66. McLaren, W., Gil, L., Hunt, S.E., Riat, H.S., Ritchie, G.R.S., Thormann, A., Flicek, P., and Cunningham, F. (2016). The Ensembl Variant Effect Predictor. Genome Biology 17, 122. 10.1186/s13059-016-0974-4.

67. Chen, S., Francioli, L.C., Goodrich, J.K., Collins, R.L., Kanai, M., Wang, Q., Alfoldi, J., Watts, N.A., Vittal, C., Gauthier, L.D., et al. (2024). A genomic mutational constraint map using variation in 76,156 human genomes. Nature 625, 92–100. 10.1038/s41586-023-06045-0.

68. Chang, C.C., Chow, C.C., Tellier, L.C., Vattikuti, S., Purcell, S.M., and Lee, J.J. (2015). Second-generation PLINK: rising to the challenge of larger and richer datasets. Gigascience 4, s13742–13015-10047-13748.

69. Thurman, R.E., Rynes, E., Humbert, R., Vierstra, J., Maurano, M.T., Haugen, E., Sheffield, N.C., Stergachis, A.B., Wang, H., Vernot, B., et al. (2012). The accessible chromatin landscape of the human genome. Nature 489, 75–82. 10.1038/nature11232.

70. Zou, H., and Hastie, T. (2005). Regularization and Variable Selection Via the Elastic Net. Journal of the Royal Statistical Society Series B: Statistical Methodology 67, 301–320. 10.1111/j.1467-9868.2005.00503.x.

71. Singh, S., Adelizzi, E., Heffner, C., Curtis, S., Duncan, K., Awotoye, W., Olotu, J., Tamara, B., Adeyemo, W., Gowans, L., et al. (2026). Zfp750 prevents oral adhesions and promotes temporary epithelial fusions. BioRxiv. 10.64898/2026.02.12.705205.

72. Farrell, J.A., Wang, Y., Riesenfeld, S.J., Shekhar, K., Regev, A., and Schier, A.F. (2018). Single-Cell Reconstruction of Developmental Trajectories during Zebrafish Embryogenesis. Science 360, 6392. 10.1126/science.aar3131.

73. Liu, H., Duncan, K., Helverson, A., Kumari, P., Mumm, C., Xiao, Y., Carlson, J.C., Darbellay, F., Visel, A., Leslie, E., et al. (2020). Analysis of zebrafish periderm enhancers facilitates identification of a regulatory variant near human KRT8/18. eLife 9, e51325. 10.7554/eLife.51325.

74. Botti, E., Spallone, G., and Moretti, F. (2011). Developmental Factor IRF6 Exhibits Tumor Suppressor Activity in Squamous Cell Carcinomas. Proceedings of the National Academy of Sciences 108, 13710–13715. 10.1073/pnas.1110931108.

75. Lopez-Pajares, V., Bhaduri, A., Zhao, Y., Gowrishankar, G., Donohue, L.K.H., Guo, M.G., Siprashvili, Z., Miao, W., Nguyen, D.T., Yang, X., et al. (2025). Glucose modulates IRF6 transcription factor dimerization to enable epidermal differentiation. Cell Stem Cell 32, 795–810 e710. 10.1016/j.stem.2025.02.017.

76. 76. Bogdanovic, O., Fernandez-Minan, A., Tena, J.J., de la Calle-Mustienes, E., Hidalgo, C., van Kruysbergen, I., van Heeringen, S.J., Veenstra, G.J., and Gomez-Skarmeta, J.L. (2012). Dynamics of enhancer chromatin signatures mark the transition from pluripotency to cell specification during embryogenesis. Genome Res 22, 2043–2053. 10.1101/gr.134833.111.

77. Khouri-Farah, N., Manchel, A., Wentworth Winchester, E., Schilder, B.M., Robinson, K., Curtis, S.W., Skene, N.G., Leslie-Clarkson, E.J., and Cotney, J. (2026). Gene expression dynamics of human and mouse craniofacial development at the single-cell level. Nat Commun. 10.1038/s41467-026-70232-6.

78. 78. Liu, J., Castillo-Hair, S.M., and Du, L.Y. (2024). Dissecting the Regulatory Logic of Specification and Differentiation during Vertebrate Embryogenesis.

79. Dworkin, S., Simkin, J., and Darido, C. (2014). Grainyhead-like 3 Regulation of Endothelin-1 in the Pharyngeal Endoderm Is Critical for Growth and Development of the Craniofacial Skeleton. Mechanisms of Development 133. 10.1016/j.mod.2014.05.005.

80. 80. de la Garza, G., Schleiffarth, J.R., Dunnwald, M., Mankad, A., Weirather, J.L., Bonde, G., Butcher, S., Mansour, T.A., Kousa, Y.A., Fukazawa, C.F., et al. (2013). Interferon Regulatory Factor 6 Promotes Differentiation of the Periderm by Activating Expression of Grainyhead-Like 3. Journal of Investigative Dermatology 133, 68–77. 10.1038/jid.2012.269.

81. Phatak, M., Kulkarni, S., Miles, L.B., Anjum, N., Dworkin, S., and Sonawane, M. (2021). Grhl3 Promotes Retention of Epidermal Cells under Endocytic Stress to Maintain Epidermal Architecture in Zebrafish. PLOS Genetics 17. 10.1371/journal.pgen.1009823.

82. Schubach, M., Maass, T., Nazaretyan, L., Roner, S., and Kircher, M. (2024). CADD v1.7: using protein language models, regulatory CNNs and other nucleotide-level scores to improve genome-wide variant predictions. Nucleic Acids Res 52, D1143–D1154. 10.1093/nar/gkad989.

83. Kotkamp, K., Mössner, R., Allen, A., Onichtchouk, D., and Driever, W. (2014). A Pou5f1/Oct4 Dependent Klf2a, Klf2b, and Klf17 Regulatory Sub-Network Contributes to EVL and Ectoderm Development during Zebrafish Embryogenesis. Developmental Biology 385, 2. 10.1016/j.ydbio.2013.10.025.

84. Liu, H., Leslie, E.J., and Jia, Z. (2016). Irf6 Directly Regulates Klf17 in Zebrafish Periderm and Klf4 in Murine Oral Epithelium, and Dominant-Negative KLF4 Variants Are Present in Patients with Cleft Lip and Palate. Human Molecular Genetics 25, 4. 10.1093/hmg/ddv614.

85. 85. Bravo Gonzalez-Blas, C., De Winter, S., Hulselmans, G., Hecker, N., Matetovici, I., Christiaens, V., Poovathingal, S., Wouters, J., Aibar, S., and Aerts, S. (2023). SCENIC+: single-cell multiomic inference of enhancers and gene regulatory networks. Nat Methods 20, 1355–1367. 10.1038/s41592-023-01938-4.

86. Howe, K., Clark, M.D., and Torroja, C.F. (2013). The Zebrafish Reference Genome Sequence and Its Relationship to the Human Genome. Nature 496, 7446. 10.1038/nature12111.

87. Hunt, W., Rebecca, A.M., Peso, L., and Wasserman, W.W. (2014). Improving Analysis of Transcription Factor Binding Sites within ChIP-Seq Data Based on Topological Motif Enrichment. BMC Genomics 15, 472. 10.1186/1471-2164-15-472.

88. Avsec, Z., Latysheva, N., Cheng, J., Novati, G., Taylor, K.R., Ward, T., Bycroft, C., Nicolaisen, L., Arvaniti, E., Pan, J., et al. (2026). Advancing regulatory variant effect prediction with AlphaGenome. Nature 649, 1206–1218. 10.1038/s41586-025-10014-0.

89. Fiers, M.W.E.J., Minnoye, L., Aibar, S., González-Blas, C.B., Atak, Z.K., and Aerts, S. (2018). Mapping Gene Regulatory Networks from Single-Cell Omics Data. Briefings in Functional Genomics 17, 246–254. 10.1093/bfgp/elx046.

90. Mangold, E., Bohmer, A.C., Ishorst, N., Hoebel, A.K., Gultepe, P., Schuenke, H., Klamt, J., Hofmann, A., Golz, L., Raff, R., et al. (2016). Sequencing the GRHL3 Coding Region Reveals Rare Truncating Mutations and a Common Susceptibility Variant for Nonsyndromic Cleft Palate. Am J Hum Genet 98, 755–762. 10.1016/j.ajhg.2016.02.013.

91. Eshete, M.A., Liu, H., Li, M., Adeyemo, W.L., Gowans, L.J.J., Mossey, P.A., Busch, T., Deressa, W., Donkor, P., Olaitan, P.B., et al. (2018). Loss-of-Function GRHL3 Variants Detected in African Patients with Isolated Cleft Palate. J Dent Res 97, 41–48. 10.1177/0022034517729819.

92. Ali, A., Bhatti, M.Z., Shah, A.S., Duong, H.Q., Alkreathy, H.M., Mohammad, S.F., Khan, R.A., and Ahmad, A. (2015). Tumor-suppressive p53 Signaling Empowers Metastatic Inhibitor KLF17-dependent Transcription to Overcome Tumorigenesis in Non-small Cell Lung Cancer. J Biol Chem 290, 21336–21351. 10.1074/jbc.M114.635730.

93. Ali, A., Zhang, P., Liangfang, Y., Wenshe, S., Wang, H., Lin, X., Dai, Y., Feng, X.H., Moses, R., Wang, D., et al. (2015). KLF17 empowers TGF-beta/Smad signaling by targeting Smad3-dependent pathway to suppress tumor growth and metastasis during cancer progression. Cell Death Dis 6, e1681. 10.1038/cddis.2015.48.

94. Gumireddy, K., Li, A., Gimotty, P.A., Klein-Szanto, A.J., Showe, L.C., Katsaros, D., Coukos, G., Zhang, L., and Huang, Q. (2009). KLF17 is a negative regulator of epithelial-mesenchymal transition and metastasis in breast cancer. Nat Cell Biol 11, 1297–1304. 10.1038/ncb1974.

95. Zhou, S., Tang, X., and Tang, F. (2016). Kruppel-like factor 17, a novel tumor suppressor: its low expression is involved in cancer metastasis. Tumour Biol 37, 1505–1513. 10.1007/s13277-015-4588-3.

96. Lea, R.A., McCarthy, A., Boeing, S., Fallesen, T., Elder, K., Snell, P., Christie, L., Adkins, S., Shaikly, V., Taranissi, M., and Niakan, K.K. (2021). KLF17 promotes human naive pluripotency but is not required for its establishment. Development 148. 10.1242/dev.199378.

97. Li, J., Tiwari, M., Xu, X., Chen, Y., Tamayo, P., and Sen, G.L. (2020). TEAD1 and TEAD3 Play Redundant Roles in the Regulation of Human Epidermal Proliferation. J Invest Dermatol 140, 2081–2084 e2084. 10.1016/j.jid.2020.01.029.

98. Wang, Z., Yang, C., Ma, Z., He, X., Li, L., Wang, Y., Wang, Y., Cai, X., Jiao, H., Zhang, M., et al. (2026). YAP-TEAD regulates the super-enhancer network to control early surface ectoderm commitment. Nucleic Acids Res 54. 10.1093/nar/gkaf1285.

99. Foreman, J., Perrett, D., Mazaika, E., Hunt, S.E., Ware, J.S., and Firth, H.V. (2023). DECIPHER: Improving Genetic Diagnosis Through Dynamic Integration of Genomic and Clinical Data. Annu Rev Genomics Hum Genet 24, 151–176. 10.1146/annurev-genom-102822-100509.

