## Supplementary figures and images for "Conservation of transcriptional regulatory networks in zebrafish and human periderm facilitates identification of *GRHL1* as an orofacial cleft risk gene"

### Supplemental Information

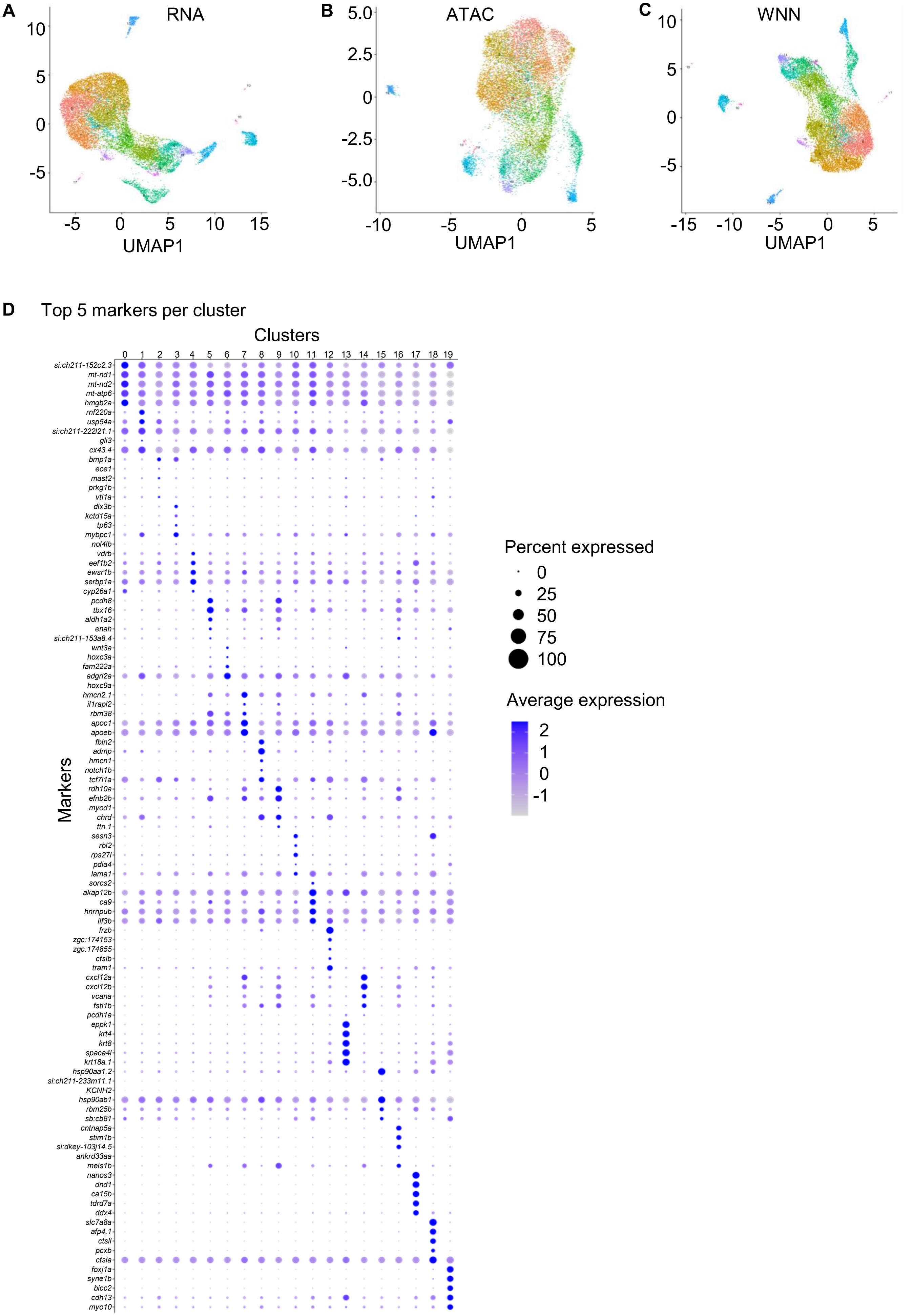

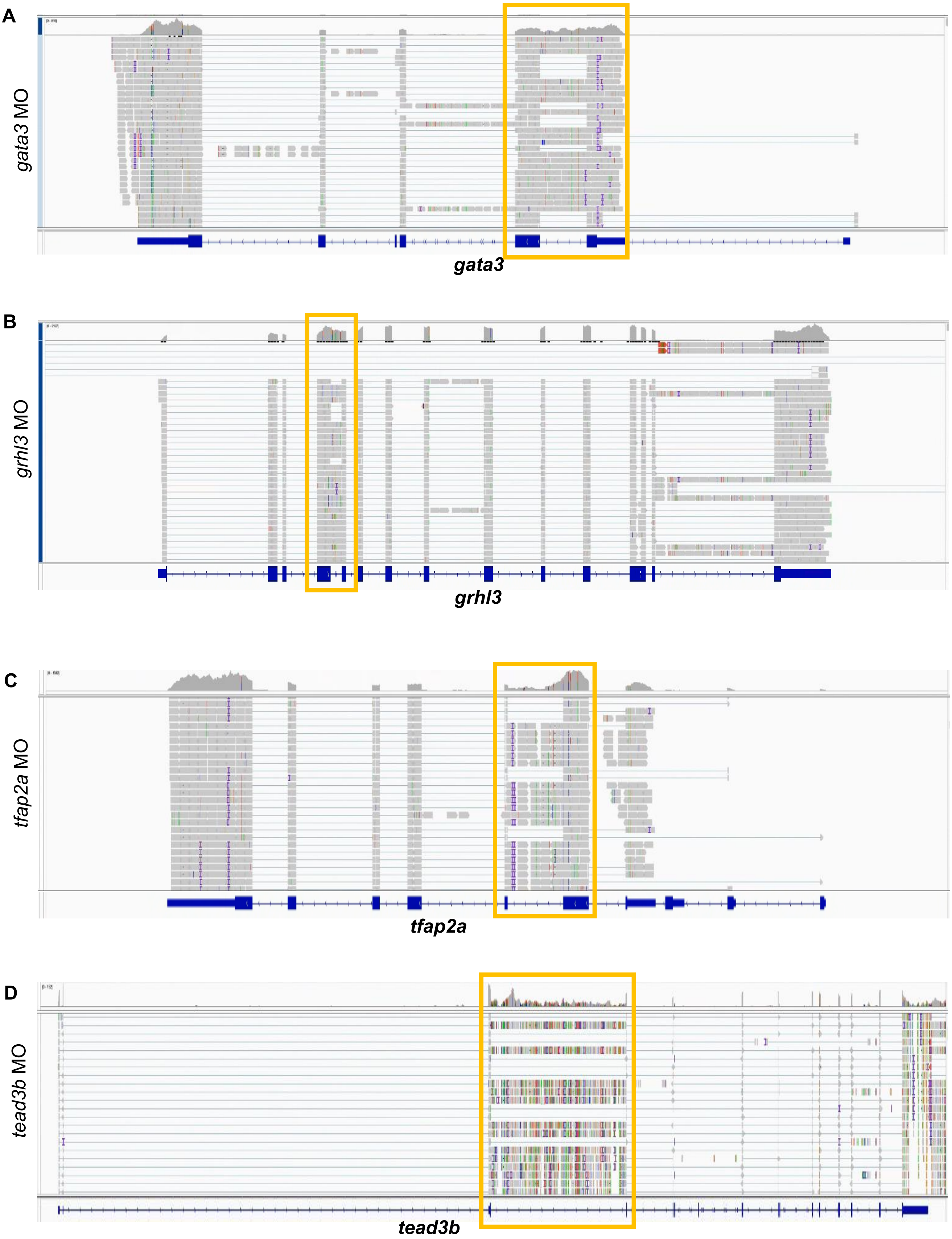

**Figure S2**

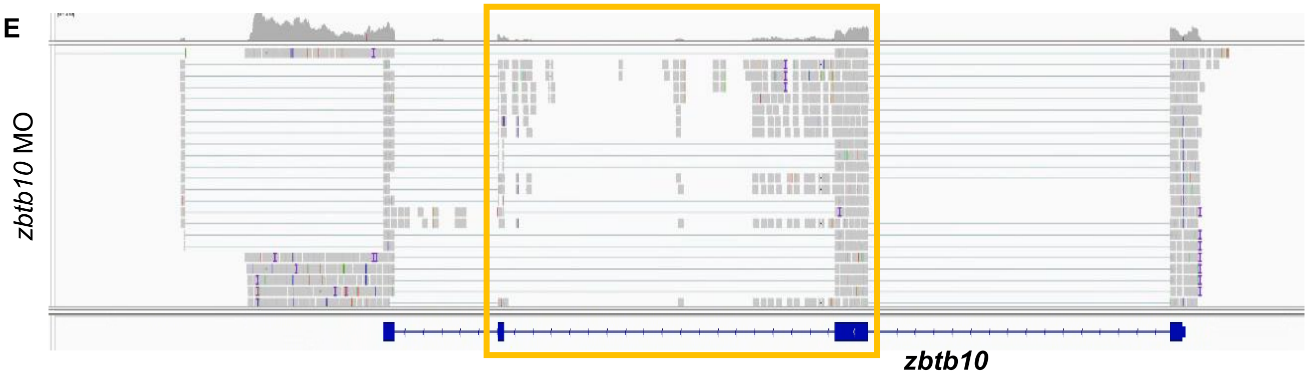

**Figure S3**

A Irf6 peaks (ChIP-seq)

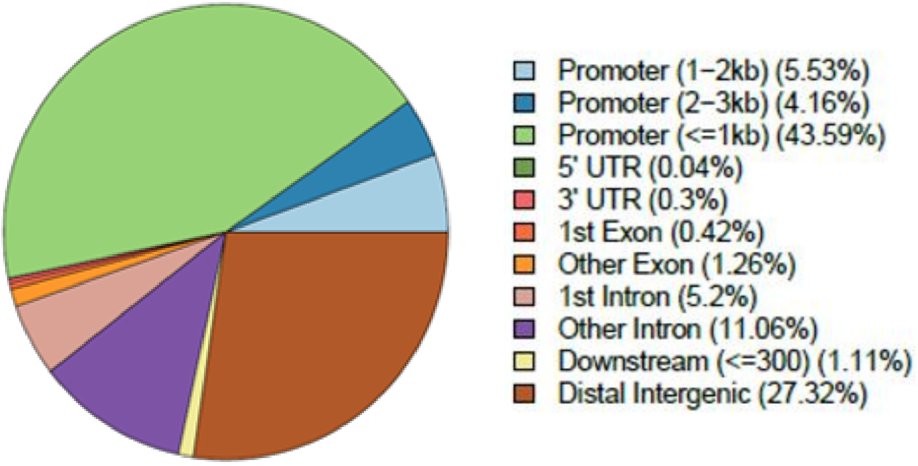


B Grhl3 peaks (CUT&RUN)

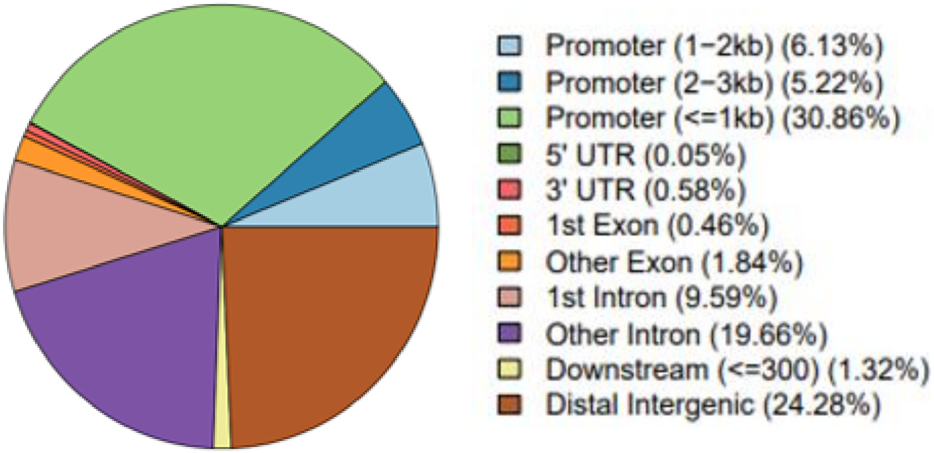


C Tfap2a peaks (CUT&RUN)

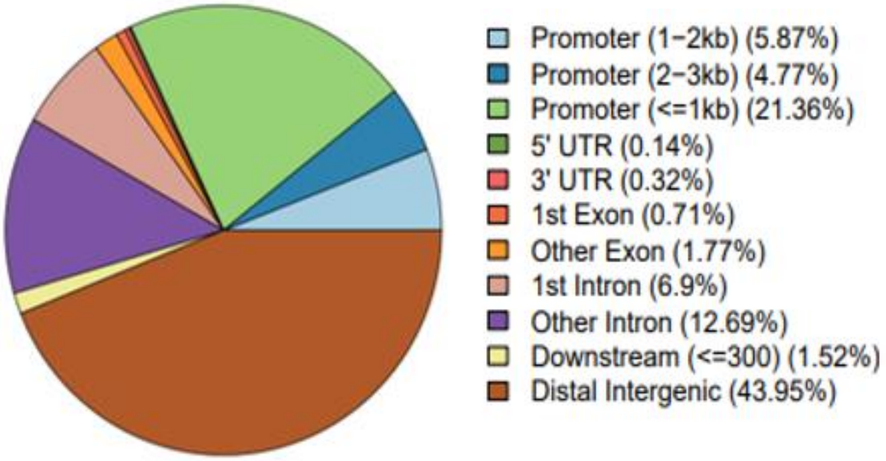

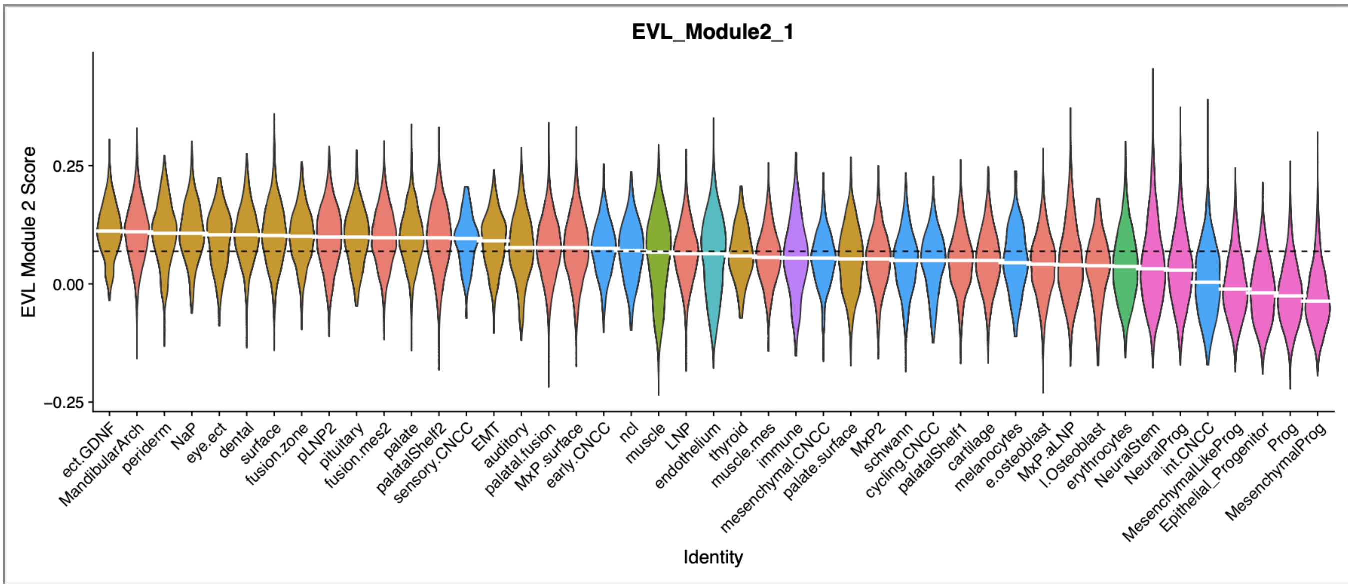

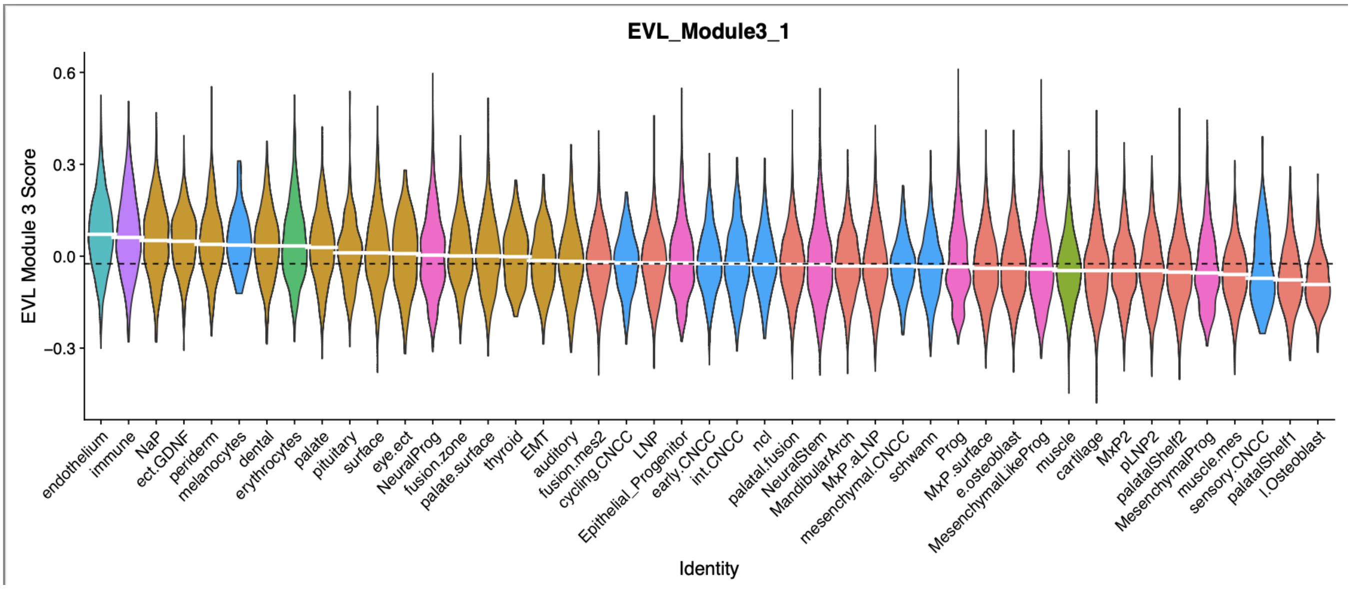

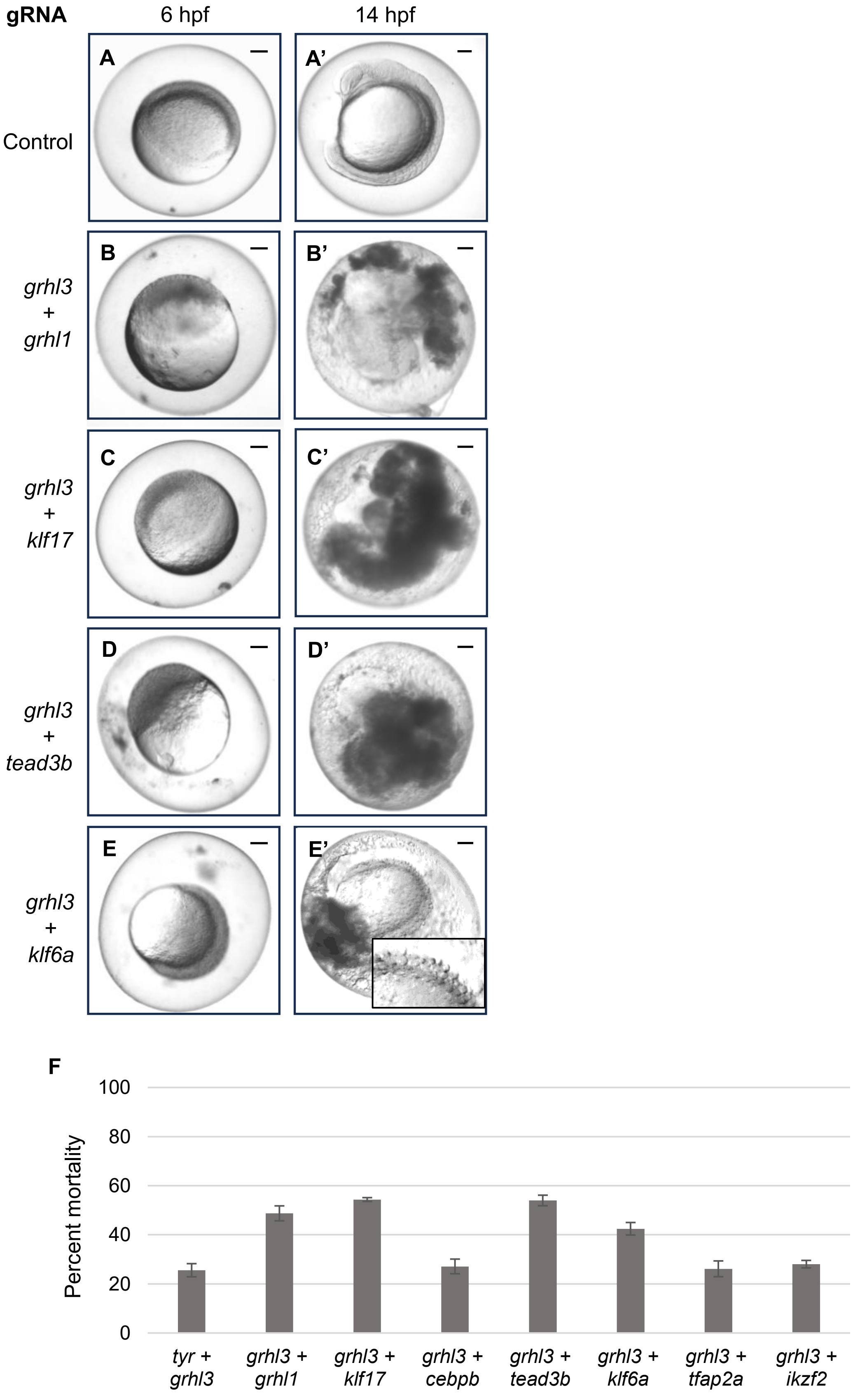

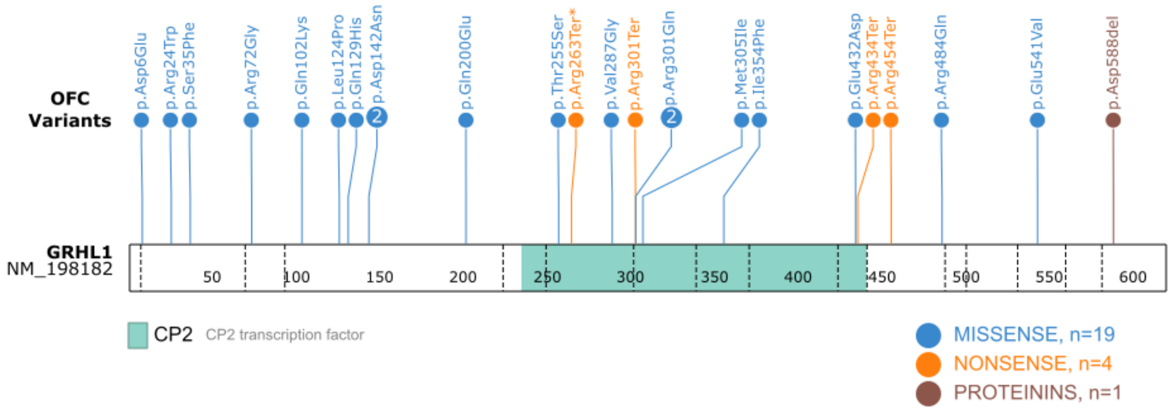
